# Characterisation of the Tilapia Lake Virus proteome and identification of an 11th protein, S9-F3

**DOI:** 10.1101/2025.09.17.676819

**Authors:** Nunticha Pankaew, Dominic Kurian, Federico De Angelis, Jorge del-Pozo, Ross D. Houston, Remi L. Gratacap, Rute M. Pinto, Paul Digard

**Affiliations:** The Roslin Institute and Royal (Dick) School of Veterinary Studies, The University of Edinburgh, Easter Bush campus, EH25 9RG, UK

**Keywords:** Tilapia, TiLV, nuclear export, accessory protein, CRM1, Leptomycin B

## Abstract

Tilapia Lake Virus (TiLV) is an emerging negative-sense, single-stranded RNA virus that poses a significant threat to global tilapia aquaculture. Of the proteins encoded by its ten genomic segments, the first four, which encode a viral polymerase (segments 1–3) and a nucleoprotein (segment 4) have been characterised. However, the functions of the polypeptides encoded by the remaining segments remain largely unknown. Here, we systematically investigated the expression and subcellular localisation of all ten predicted TiLV-encoded proteins using *in vitro* translation and expression in mammalian and fish cells as well as mass spectrometry of virus infected cells. These approaches confirmed the synthesis of major polypeptides from each segment and identified an additional 11th protein, S9-F3, translated from an alternative reading frame of segment 9. Microscopy of individually expressed green fluorescent protein (GFP)-tagged constructs revealed that S2 and S10 polypeptides were predominantly nuclear, while S1, S3, S5, S8 and S9-F3 were mostly cytoplasmic with S5 and S6 displaying perinuclear foci. Bioinformatic analysis suggested a potential nuclear export signal in S9-F3 and use of the inhibitor leptomycin B confirmed its CRM1-dependent nuclear export. Evolutionary analyses indicated that both S9 and S9-F3 are under selective pressure, as well as the presence of an S9-F3 homologue in a TiLV-like guppy virus. These findings uncover an alternative translation product and a regulated nuclear export mechanism in TiLV, providing new insights into the molecular biology of this virus and its interaction with host cellular pathways.

## Introduction

Tilapia lake virus (TiLV) is an emerging orthomyxo-like virus that poses a major threat to global tilapia aquaculture. First reported in 2014, TiLV has been associated with mortality rates of up to 90% in affected fish populations (Bacharach et al., 2014). Since its discovery, the virus has been detected across multiple continents, including Asia, Africa, and North America (Aich et al., 2022), indicating the urgent need for improved surveillance and disease management strategies. Given that Nile tilapia *(Oreochromis niloticus*) ranks among the top aquaculture species worldwide in terms of production (FAO, 2024), TiLV represents a serious constraint to global food security and aquaculture sustainability.

TiLV is classified as *Tilapia tilapinevirus* within the newly established family Amnoonviridae, under the order Articulavirales, which also includes influenza viruses (Adams et al., 2017). Its genome comprises 10 negative-sense, single-stranded RNA segments, each bearing conserved, quasi-complementary termini flanking an open reading frame (ORF), like other segmented RNA viruses (Bacharach et al., 2016). The primary sequences of the putative viral proteins encoded by the TiLV segments mostly lack homology to known proteins, with the exception of segment 1, which contains conserved motifs characteristic of an orthomyxovirus PB1-like polymerase subunit (Bacharach et al., 2016). This lack of sequence homology has hindered straightforward characterisation of the remaining viral proteins. To date, only the first four segments have been functionally annotated: structural studies have shown that segments 1–3 encode the viral RNA-dependent RNA polymerase (Arragain et al., 2023), while biochemical and structural analyses identify segment 4 as encoding the nucleoprotein (NP) (Abu Rass et al., 2022; Arragain et al., 2025). The functions of proteins encoded by the remaining six segments remain largely undefined, and the possibility of further accessory proteins being expressed via alternative translation mechanisms, as seen in orthomyxoviruses (Pinto et al., 2021) remains unexplored.

In RNA viruses, alternative translation strategies such as leaky ribosomal scanning and overlapping open reading frames (ORFs) enable the expression of multiple proteins from a single mRNA (Firth & Brierley, 2012; Pinto et al., 2021). Translation initiation in eukaryotes is regulated in part by the Kozak consensus sequence surrounding the AUG start codon, which influences ribosome recognition and the likelihood of translation initiation (Kozak, 1986). Suboptimal Kozak contexts at the first AUG can lead to leaky ribosomal scanning, allowing downstream AUGs to initiate translation and potentially producing additional polypeptides. This mechanism has been well-documented in influenza A virus (IAV), where accessory proteins such as PB1-F2 and PB1-N40 arise from alternative AUG usage within the same mRNA to either access another reading frame or to produce an N-terminally truncated derivative of the canonical segment polypeptide (Chen et al., 2001; Jaafar & Kieft, 2019; Wise et al., 2009). Alternatively, programmed ribosomal frameshifting events can produce a polypeptide initiated from the canonical transcript AUG codon but with a divergent C-terminal domain, such as IAV PA-X (Jagger et al., 2012). These proteins can modulate host immune responses, enhance viral replication, and contribute to pathogenesis (Cheung et al., 2020; Hu et al., 2018). Diversification of viral proteomes by such mechanisms is seen as an evolutionary response to the size constraints imposed by an RNA genome (Belshaw et al., 2008). The compressed nature of the TiLV genome (10.3 kb in total with the largest segment only 1641 nucleotides long (Bacharach et al., 2014)); suggests it too might express alternative polypeptides.

In this study, we systematically characterised the coding capacity of TiLV by expressing individual TiLV segments *in vitro* in both a cell-free system and in transfected cells, to examine their expression capacity and, using GFP-tagged versions, the subcellular localisation of TiLV proteins. Viral protein expression was confirmed by mass spectrometry of virus-infected fish cells. These approaches allowed us to show that at least one polypeptide species is expressed from each TiLV segment and led to the discovery of an 11th TiLV protein encoded by an ORF embedded in an alternative reading frame of segment 9.

## Results

### Prediction of functional motifs in TiLV polypeptides

To predict functional motifs and any potential accessory proteins encoded by TiLV proteins, all ten viral gene segments were analysed using computational biology approaches. The major ORF within each segment was identified across the three plus-strand reading frames, and AUG codons within the canonical ORF, along with any alternative downstream AUG codons in frames 2 and 3 ahead of a ≥ 25 codon ORF, were identified and plotted (Figure 1) to assess the potential for alternative translation initiation events. To provide further context, the diagrams were further annotated with protein domain information for the five TiLV polypeptides with solved crystal structures (Arragain et al., 2025; Arragain et al., 2023; Wang, 2024). Examination of the sequences surrounding the primary AUG codon of each segment suggested that the canonical AUG codons of S3, S4, and S8 ORFs are situated within a strong Kozak context (Table S1), assuming that the translation initiation apparatus of tilapia follows the conserved mammalian-like pattern of zebrafish (Grzegorski et al., 2014). In contrast, the primary AUGs of S1, S2, S7, S9, and S10 ORFs have an intermediate strength Kozak consensus while those of S5 and S6 have weak motifs, indicating the potential for leaky ribosomal scanning events. For most of these segments, there are no major ORFs with initiator AUG codons in frames 2 or 3 near the 5’-end of the segment that could be accessed by a scanning ribosome that had missed the main ORF AUG (Figure 1), leaving the possibility of leaky scanning producing N-terminal variants of the primary protein. The exception, as previously noted (Acharya et al., 2019), is segment 9, which contains a large 105 codon alternative ORF in frame 3, only 11 codons shorter than the primary S9 ORF. This trans-frame ORF, (denoted as S9-F3 from hereon), possesses an AUG in a strong Kozak consensus, supporting the possibility of its expression via leaky ribosomal scanning. Finally, we considered whether TiLV might possess a PA-X analogue. In IAV segment 3, the PA-X protein is produced by a +1 ribosomal frameshift event to access the X ORF, which almost exactly overlaps the coding region of the linker region between N-terminal endonuclease and C-terminal domains of PA (Firth et al., 2012; Shi et al., 2012). However, while TiLV S3 has two +1 ORFs of 41 and 39 codons, broken by an intervening stop codon, that overlap the linker coding region in S3 (Figure 1), we could not identify an obvious PA-X-like frameshift signal.

**Figure 1.**
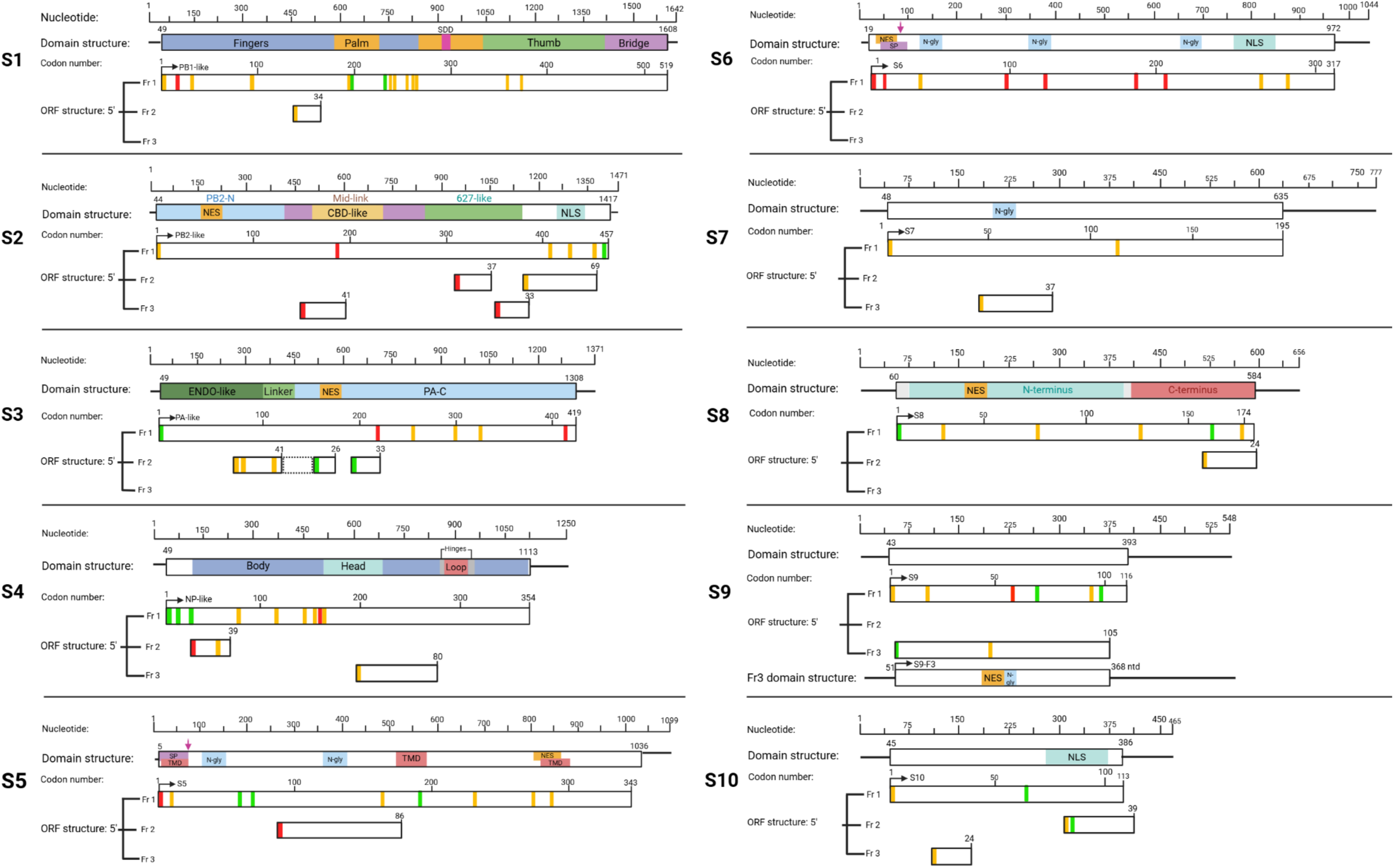
Schematic representation of the ORFs and predicted sequence motifs of TiLV segments 1–10. UTRs are symbolised by horizontal black lines, and open boxes represent ORFs, approximately to scale. The primary coding regions are annotated (top bars) with predicted functional motifs (SDD, RdRp catalytic motives (fingers, palm and thumb); NLS, nuclear localisation signal; SP, signal peptide; N-gly, N-linked glycosylation; TMD, transmembrane domain; NES, nuclear export signal). Structural domains of S1-S4, corresponding to the influenza polymerase subunits and NP respectively, and S8 were taken from PDB 8PSN, 9HBR and 8IXW, respectively. Other ORFs (minimum length 25 codons) starting with a methionine codon from the three frames (Fr) are also indicated. The box with dashed lines in S3 represents an ORF lacking AUG codons in the region corresponding to the X-ORF in IAV. AUG codons represented by vertical lines are coloured according to the predicted strength of their Kozak translation initiation context (green, strong; amber, intermediate; red, weak). The predicted major polypeptides are shown in frame 1 (Fr1) with black arrows indicating canonical AUGs predicted to be used for translation initiation of the primary TiLV protein.

Previous sequence analysis of S1 (Bacharach et al., 2016) revealed a catalytic RNA-dependent RNA polymerase (RdRp) SDD motif at amino acids 288–290 (Figure 1, Table S2), located within the palm sub-domain of the viral RdRp (Arragain et al., 2023). To identify additional sequence motifs indicative of potential functions in other TiLV proteins, we used a suite of online tools: cNLS Mapper (Kosugi et al., 2008), SignalP (Almagro Armenteros et al., 2019), NetNGlyc (Gupta & Brunak, 2002), TMHMM (Krogh et al., 2001; Sonnhammer et al., 1998) and LocNES (Xu et al., 2015). These computational analyses predicted nuclear localisation signal (NLS) and/or nuclear export signals (NES) in S2, S3, S5, S6, S8, S9-F3 and S10 ORFs, suggesting potential nucleocytoplasmic transport. While no functional motifs were identified in the primary ORFs of segments 4 and 9, multiple ones were predicted in the S5 and S6 proteins, including a cleavable signal peptide (SP), N-glycosylation (N-gly) sites, NES, and transmembrane domains (TMDs, only in S5), as well as an NLS found only in S6. These features indicate that S5 may play a role in the viral envelope structure. The predicted SP cleavage site in S5 matches a previous prediction by Bacharach et al. (2016), but the SP cleavage probability score for S6 appears relatively low (Table S2). Additionally, potential glycosylation motifs were identified within the primary ORF of S7 and the S9-F3 ORF; however, their functional relevance is uncertain due to the absence of motifs that would direct the proteins to the endoplasmic reticulum (ER).

### Expression and cellular localisation of S1–10 proteins

To determine the polypeptides translated from TiLV segments, we created a set of plasmids containing synthetic cDNAs corresponding to TiLV segments 1-10 under the control of a bacteriophage T7 RNA polymerase promoter and used these to programme *in vitro* protein synthesis in a rabbit reticulocyte lysate system, supplemented with a mix of [^35-^S] methionine and [^35-^S] cysteine. Radiolabeled protein detection successfully identified polypeptides translated from all ten segments, each yielding a major product that generally corresponded to the predicted molecular weight of the product from the primary ORF (Figure 2A, red arrowheads). However, the S10 product consistently migrated unexpectedly slowly on SDS-PAGE, at an apparent molecular weight of approximately 18 kDa—substantially higher than its predicted size of ∼13 kDa. In addition to the major bands corresponding to the predicted protein sizes, several prominent but lower-abundance products were also detected during the translation of S1, S5, S6, and S9, consistent with the suboptimal Kozak consensus of their first AUG codon and the potential occurrence of alternative translational events.

**Figure 2.**
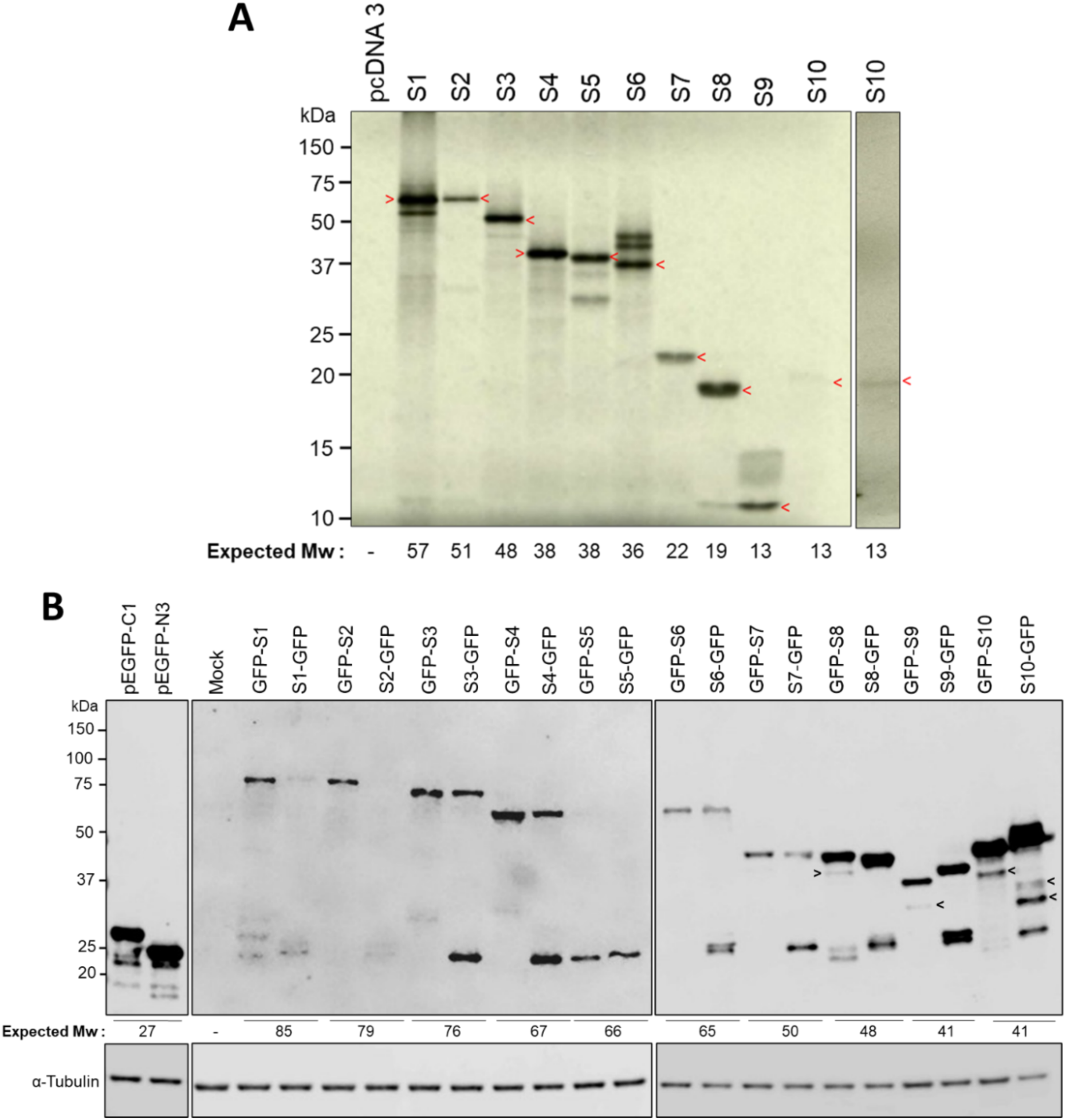
Expression of TiLV proteins *in vitro* and in cells. (A) *In vitro* translation of TiLV segments 1–10. Full-length coding sequences of each segment were transcribed and translated in rabbit reticulocyte lysate supplemented with [³⁵S]-methionine/cysteine. Lysates were resolved by SDS-PAGE and translation products visualised by autoradiography. Red arrowheads indicate the predominant protein products. An additional independent experiment (lane 13) confirmed slow migration of the S10 product. (B) Expression of GFP-tagged TiLV proteins in 293T cells. Cells were transfected with constructs encoding each polypeptide (S1–S10) fused to GFP, or with empty vectors. 48 h later, cell lysates were subjected to SDS-PAGE and immunoblotting using anti-GFP and anti-α-tubulin (loading control) antibodies.

Following successful expression of polypeptides from the ten TiLV segments *in vitro*, we next investigated their expression in readily transfectable mammalian 293T cells. At project start, specific antibodies against TiLV proteins were not available. Therefore, to enable detection in a cellular context, each primary ORF was tagged with the green fluorescent protein (GFP) gene to enable product detection through intrinsic fluorescence or via anti-GFP antibodies. To mitigate positional effects of the GFP tag, each TiLV protein was fused to GFP at either the N- or C-terminus and the resulting constructs were then transfected into both mammalian and tilapia cells. Western blot analysis using anti-GFP antibodies confirmed the expression of S1, S2, S3, S4, S6, S7, and S8 at the expected molecular weights, although constructs with GFP at the N-terminus generallly expressed to higher levels and with lower amounts of products migrating around the size (25 kDa) expected for a detached GFP "stub" (Figure 2B). S1-GFP and S2-GFP expressed particularly poorly. However, both N- and C-terminal GFP-tagged versions of S5 failed to produce detectable fusion proteins of the correct size. The mobility of GFP-tagged S9 in SDS-PAGE varied with the tag’s position, while S10 fusion proteins also showed discrepancies between N- and C-terminal tags and migrated more slowly than expected (∼50–52 kDa), consistent with the cell-free *in vitro* translation results.

Next, we transfected the set of GFP-tagged expression constructs into both mammalian 293T and tilapia OmB cells to determine the intracellular localisation of the TiLV proteins. Fluorescence imaging confirmed successful expression (including both S5 constructs, despite the failure to detect full-length products by western blotting) in both cell types (Figure S1). Higher transfection efficiency and/or stronger protein expression was consistently observed in 293T cells compared to OmB cells, perhaps reflecting compatibility issues with the mammalian RNA polymerase II promoter used for plasmid construction and/or transfection limitations. Nevertheless, both cell types gave sufficient expression to permit conclusions to be drawn over cellular compartmentalisation of the TiLV proteins, with the exception of S2-GFP, where too few fluorescent cells could be found for reliable imaging.

When imaged using higher resolution by confocal microscopy, both cell types gave negligible levels of autofluorescence in untransfected cells, while GFP alone localised diffusely throughout the cell (Figure 3). Considering the GFP-TiLV fusions first, due to the lower levels of potentially-confounding fluorescence from GFP-sized polypeptides visible by western blotting; GFP-S2 and -S10 were almost exclusively nuclear (though partially excluded from specific sub-nuclear domains, probably nucleoli) in both human and tilapia cells, consistent with the predictions of C-terminal NLS signals in both proteins. However, the other two polymerase proteins, GFP-S1 and -S3, were mostly but not entirely cytoplasmic in human cells, but while S1 was also mostly cytoplasmic in OmB cells, S3 showed a more even distribution in OmB cells. Consistent with previously reported immunofluorescent studies of TiLV-infected cells (Abu Rass et al., 2022), the other major RNP component. S4, formed large puncta in both mammalian and fish cells, although these were cytoplasmic in 293T and nuclear in OmB cells. GFP-S6, another TiLV protein with a potential NLS, localised to both nuclear and cytoplasmic compartments, though more strongly to the cytoplasm, sometimes showing perinuclear foci. Similarly, GFP-S7 and -S9 showed fluorescence signal in both cellular compartments while GFP-S8 was strongly excluded from the nucleus in human but not fish cells.

**Figure 3.**
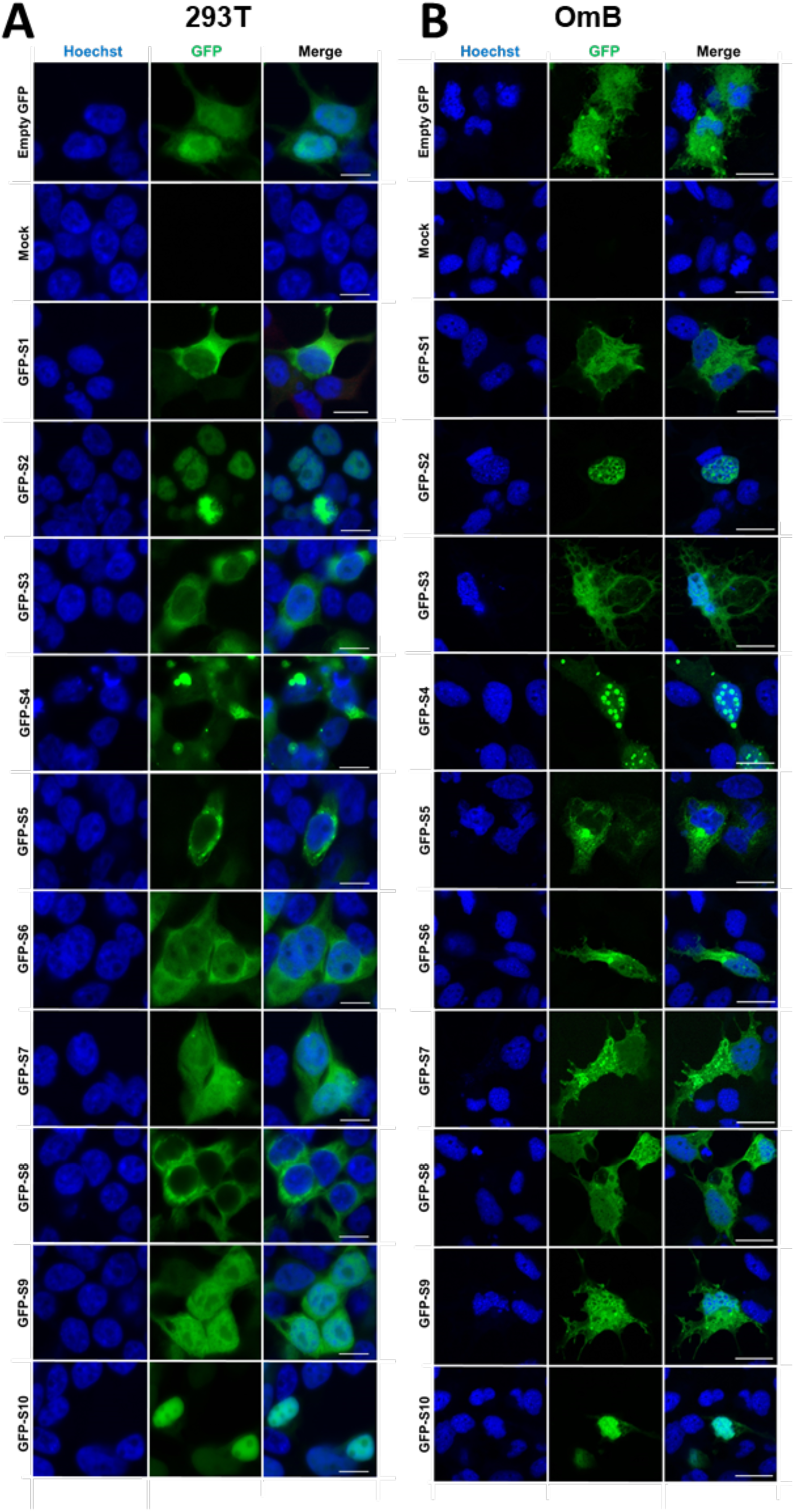
Subcellular localisation of GFP-tagged TiLV proteins in mammalian and tilapia cells. (A) 293T and (B) OmB cells were transfected with plasmids expressing N-terminal GFP fusion constructs of TiLV segments 1–10, an empty GFP vector (pEGFP-C1), or were mock-transfected. 48 h post-transfection, cells were fixed and stained with Hoechst dye to visualise nuclei. GFP (green) and Hoechst (blue) fluorescence images were acquired using a Zeiss LSM710 microscope at using an 63x oil objective. Images are single optical slices, representative of three independent experiments, each performed with a single technical replicate. Scale bars: 10 µm.

Finally, GFP-S5 was also excluded from the nucleus in both cell types, but formed perinuclear foci reminiscent of proteins targeted to the secretory pathway, consistent with the presence of SP and TM domains in its primary sequence. Examination of the C-terminally GFP tagged samples largely corroborated these observations. S10-GFP was strongly nuclear in both cell types, as was S2-GFP in 293T cells, while the other two subunits of the polymerase were predominantly cytoplasmic (Figure S2). S4-GFP was cytoplasmic in human cells and nuclear in fish cells, but produced fewer large puncta than the N-terminally tagged version. S7- and S9-GFP localised throughout the cells, while S8-GFP was strongly cytoplasmic. S5-GFP remained cytoplasmic but displayed fewer perinuclear foci than GFP-S5. Conversely, S6-GFP was more cytoplasmic than GFP-S6 and also showed a greater tendency to produce perinuclear foci. In summary, several distinct patterns of subcellular localisation for TiLV proteins were identified (Table 1), providing initial insights into their functions.

**Table 1.** Subcellular localisation of GFP-fusion TiLV proteins in different cell lines. C, cytoplasmic only; N, nuclear only; N+C, both cytoplasmic and nuclear; N_p_+C_p_, punctate; PrN, perinulear; n/d, not determined.

| Segment | Plasmid | Protein localisation |  |
| --- | --- | --- | --- |
|  |  | 293T | OmB |
| 1 | GFP-S1 | C | C |
|  | S1-GFP | C | C |
| 2 | GFP-S2 | N | N |
|  | S2-GFP | N | n/d |
| 3 | GFP-S3 | C | N+C |
|  | S3-GFP | C | N+C |
| 4 | GFP-S4 | N<sub>p</sub>+C<sub>p</sub> | N<sub>p</sub>+C<sub>p</sub> |
|  | S4-GFP | N+C | N+C |
| 5 | GFP-S5 | C, PrN | C, PrN |
|  | S5-GFP | C, PrN | C, PrN |
| 6 | GFP-S6 | C | C |
|  | S6-GFP | C | C |
| 7 | GFP-S7 | N+C | N+C |
|  | S7-GFP | N+C | N+C |
| 8 | GFP-S8 | C | N+C |
|  | S8-GFP | C | C |
| 9 | GFP-S9 | N+C | N+C |
|  | S9-GFP | N+C | N+C |
| 10 | GFP-S10 | N | N |
|  | S10-GFP | N | N |

### Mass spectrometry analysis of TiLV-infected cells

To probe the expression of TiLV-encoded proteins within a viral infection context, proteomic analysis was performed on TiLV-infected OmB cells. Cells were infected at a MOI of 0.01, and after 10-days, cell pellets were harvested and processed for MS analysis (Figure 4A). The samples underwent standard proteomic workflows: total protein extraction, quantification, in-solution digestion (reduction, alkylation, and tryptic digestion), and peptide purification. To detect both known and potential novel TiLV proteins, the resulting peptides were analysed by liquid chromatography-tandem mass spectrometry (LC-MS/MS) and searched against a custom database comprising the *Oreochromis niloticus* reference proteome (NCBI RefSeq: GCF_001858045.2) and a six-frame translation of the TiLV genome (GenBank Accession Nos. KU751814–KU751823). All ORFs (with or without start codons) ≥ 10 codons long were included (Table S3). Peptide identification was performed using standard database search algorithms, and protein quantification was achieved via label-free quantification, based on peptide precursor ion intensities and area-under-the-curve chromatographic measurements. The semi-quantitative nature of the MS analysis enabled estimation of relative protein abundance across samples. Approximately 5,000 tilapia-derived proteins were detected in both mock- and TiLV-infected samples, with abundances ranging from ∼5 × 10⁶ to ∼30 arbitrary units (a.u.) (Figure 4B and Table S4). No viral peptides were confidently detected in mock-infected samples, except for S1, which had minimal coverage (3%) and low spectral counts (Table 2), perhaps representing cross contamination. In contrast, 11 distinct TiLV-derived proteins were confidently identified in infected cells, each supported by ≥ 4 unique high-quality peptide matches (Tables 2 and S4) and substantial sequence coverage. This included the expected ten gene products encoded by the TiLV genome, as well as the non-canonical polypeptide S9-F3 translated from an alternative +2 reading frame of the S9 mRNA. All detected TiLV proteins ranked in the upper half of the protein abundance distribution (Figure 4B). Among them, S4 was the most abundant, ranking 6th overall with an abundance of 2.4 × 10⁶ a.u., followed by the three polymerase subunits at ∼ 5 x 10^5^ a.u., while the least abundant was the alternative, trans-frame product S9-F3, detected at 1.7 × 10⁴ a.u.

**Figure 4.**
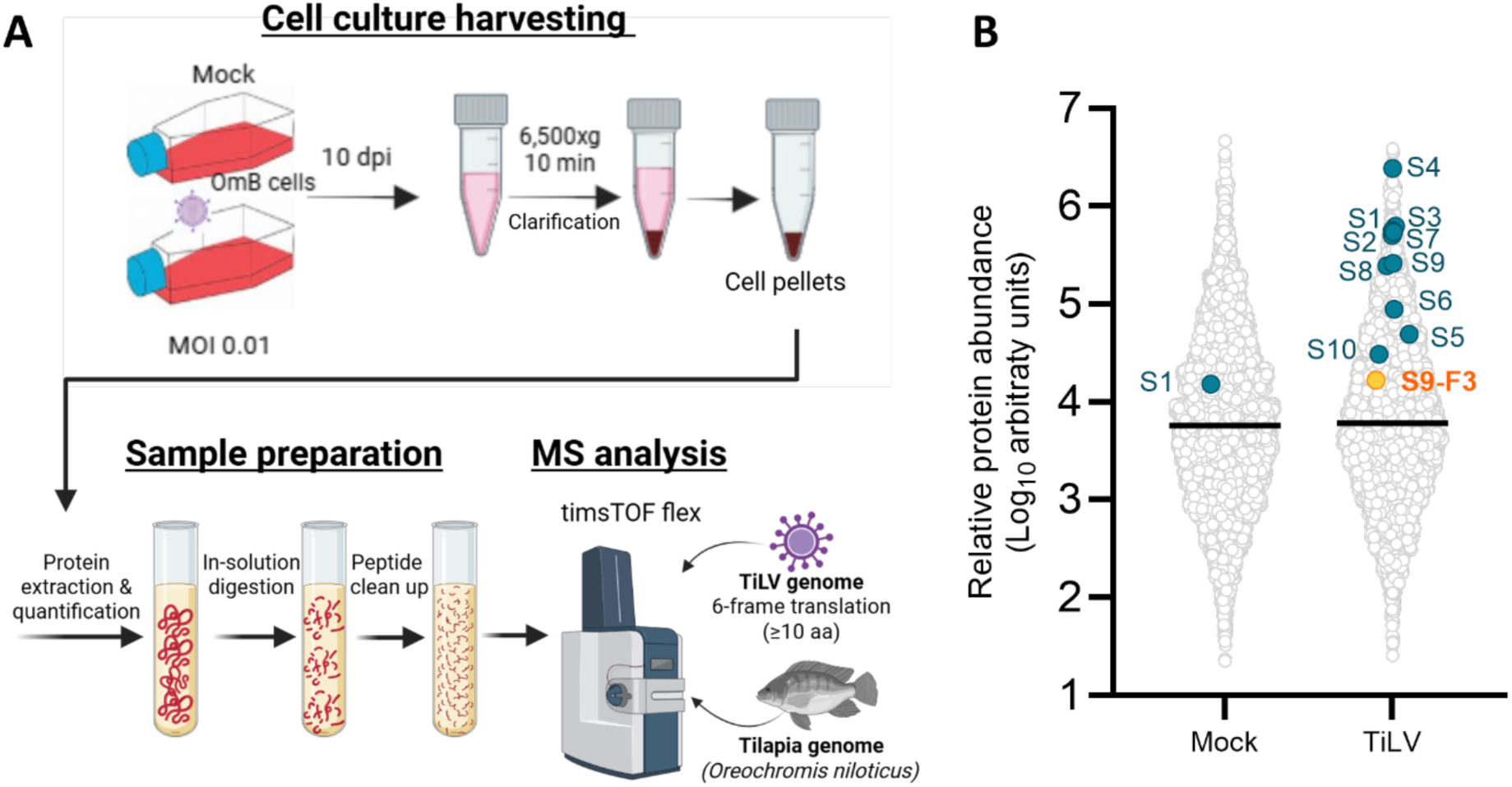
Identification of TiLV gene products from infected cells. (A) Workflow for proteomic analysis of TiLV-infected cells. OmB cells infected or mock-infected with TiLV (MOI 0.01) were harvested at 10 dpi, and cell pellets were subjected to sample preparation and mass spectrometry (MS) analyses, cross-referencing against tilapia (*Oreochromis niloticus*) and TiLV (Til-4-2011 strain) genomes. (B) Proteomic identification of TiLV and cellular polypeptides in OmB cells. The relative abundance (arbitrary units) of all proteins identified by ≥ 2 unique peptides are plotted. Cellular proteins are indicated by open circles, TiLV proteins by labelled coloured dots.

**Table 2.** Mass spectrometry detection of TiLV proteins in mock and infected OmB cells. Protein molecular mass (Da), peptide identification confidence score (-10lgP), total and unique peptides (that only mapped to that protein in our database), sequence coverage (%), protein abundance (area under the curve in arbitrary units, a.u.), and MS/MS spectra counts are reported. ND = not detected.

| TiLV proteins | Mass (Da) | -10lgP | Unique peptides | Coverage (%) |  | Area (a.u.) |  | MS/MS spectra |  |
| --- | --- | --- | --- | --- | --- | --- | --- | --- | --- |
|  |  |  |  | Mock | TiLV | Mock | TiLV | Mock | TiLV |
| S1 | 57165 | 398.2 | 49 | 3 | 70 | 1.5 x 10<sup>4</sup> | 6.3 x 10<sup>5</sup> | 3 | 180 |
| S2 | 51355 | 417.8 | 42 | 0 | 74 | ND | 5.0 x 10<sup>5</sup> | 0 | 165 |
| S3 | 48185 | 373.8 | 40 | 0 | 80 | ND | 5.7 x 10<sup>5</sup> | 0 | 190 |
| S4 | 38962 | 401.2 | 54 | 0 | 71 | ND | 2.4 x 10<sup>6</sup> | 0 | 262 |
| S5 | 38186 | 223 | 7 | 0 | 35 | ND | 4.9 x 10<sup>4</sup> | 0 | 16 |
| S6 | 37084 | 282.7 | 18 | 0 | 51 | ND | 8.9 x 10<sup>4</sup> | 0 | 49 |
| S7 | 23621 | 306.1 | 30 | 0 | 67 | ND | 5.5 x 10<sup>5</sup> | 0 | 97 |
| S8 | 21723 | 235.1 | 11 | 0 | 27 | ND | 2.5 x 10<sup>5</sup> | 0 | 40 |
| S9 | 15009 | 195.6 | 5 | 0 | 37 | ND | 2.6 x 10<sup>5</sup> | 0 | 44 |
| S9-F3 | 13696 | 142.2 | 4 | 0 | 49 | ND | 1.7 x 10<sup>4</sup> | 0 | 9 |
| S10 | 12959 | 108.3 | 4 | 0 | 18 | ND | 3.1 x 10<sup>4</sup> | 0 | 11 |

### Functional conservation and evolutionary analysis of S9 and S9-F3

The discovery of S9-F3 prompted further investigation into its evolutionary relationship with the primary S9 protein. To investigate the temporal evolutionary dynamics of segment 9 and its overlapping gene within the TiLV population, a time-based phylogenetic reconstruction was performed for a dataset of aligned segment 9 nucleotide sequences using Bayesian Evolutionary Analysis Sampling Trees (BEAST) (Suchard et al., 2018). Our dataset (Table S5) of publicly-available sequences included 44 TiLV sequences spanning nine geographically diverse outbreak regions and a variety of host species, as well as two TiLV-like viruses sequenced from guppy fish *(Poecilia reticulata)* (Li, 2023). A further unnamed TiLV sequence lacking metadata (MW281464.1) was excluded from the BEAST analysis but included in downstream analyses. Metadata on host species and geographic origin were incorporated as discrete traits, and a maximum clade credibility (MCC) tree was summarised in R. The resulting phylogenetic tree (Figure 5A) suggests that the last common ancestor of segment 9 from currently sampled TiLV isolates was in the early 1980s and that these diverged from the guppy TiLV-like virus around a century ago. Asia, and within that continent, Thailand, contributed the highest number of sequences, followed by India and Bangladesh, reflecting both disease burden and surveillance intensity. However, no particular phylogeographic pattern of segment 9 evolution was apparent. Most sequences originated from *Oreochromis niloticus*, but additional hosts such as *O. aureus* and *Sarotherodon galilaeus* were also represented, implying a broad host range without clear evidence of host-restricted evolution.

**Figure 5.**
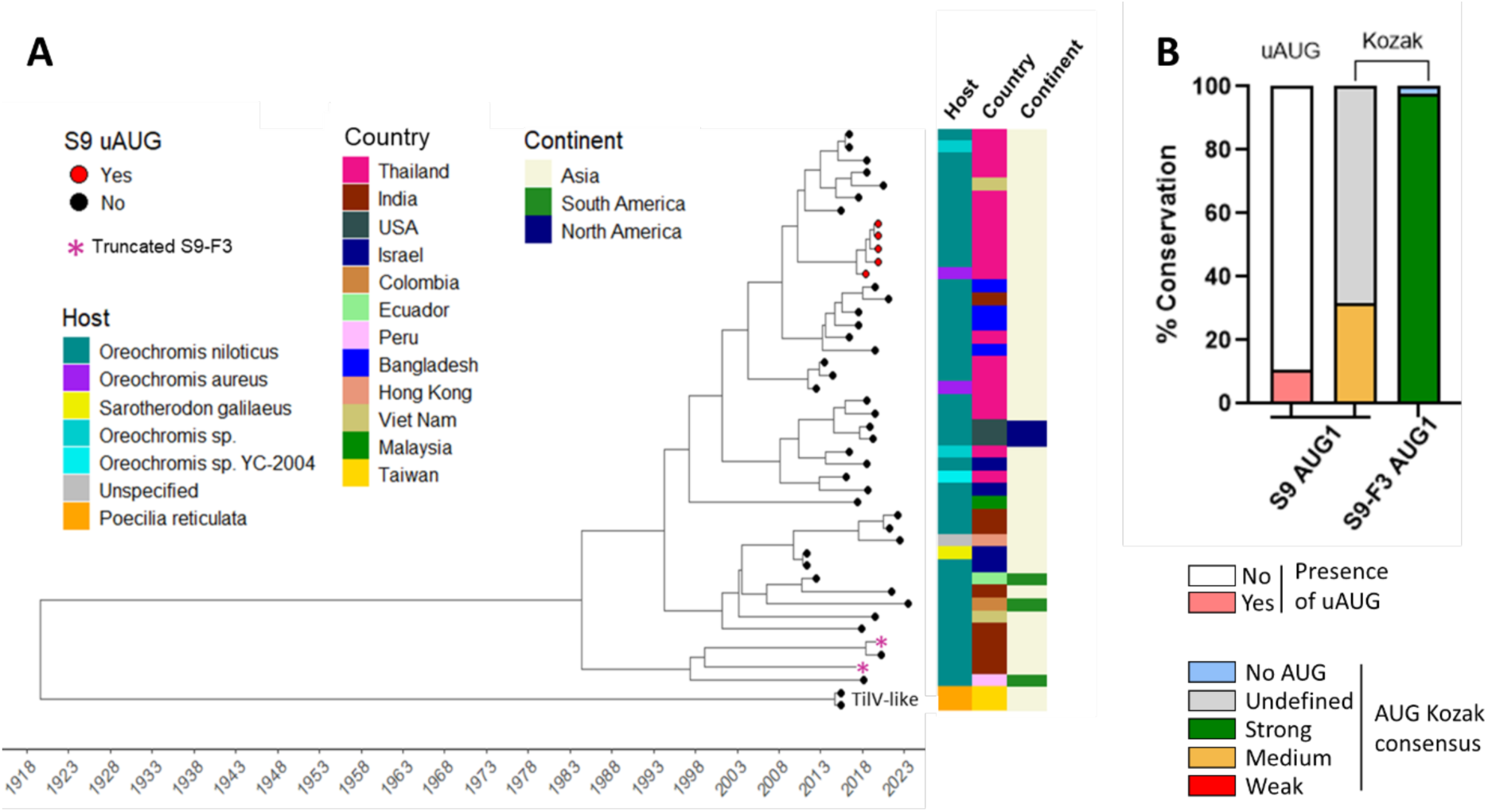
Phylogenetic tree of TiLV Segment 9 and conservation of 5’ initiator AUGs. (A) Time-scaled phylogenetic tree of TiLV and TiLV-like segment 9 sequences (n = 46). Phylogeny was inferred using BEAST. Tip points are coloured according to the presence (red) or absence (black) of an upstream AUG in the S9 ORF. Asterisks denote sequences with a truncated S9-F3 ORF. Bars to the right annotate host species, country, and continent of origin. (B) Analysis of translation initiation context for TiLV S9 and S9-F3. Segment 9 sequences were aligned and the Kozak consensus sequence surrounding potential AUG initiation codons for S9 and S9-F3 were examined and color-coded according to predicted translation initiation strength as previously described. Codons with undefined Kozak consensus are shown in grey, and those lacking an AUG are in light blue. No segment with an uAUG had sufficient UTR sequence reported to define a Kozak consensus.

To investigate S9-F3 conservation, we examined the ORF structures of the segment 9 dataset. All TiLV and TilV-like segment 9 sequences contain the canonical S9 AUG codon, mostly as the initiator of a 116-codon ORF (Figures S3, S4A). However, a distinct monophyletic cluster of five Thai isolates from circa 2018 (Figures 5A, S3) share a novel upstream start codon (uAUG) in the segment UTR, which, if used, would produce a 119 amino-acid S9 protein, while one isolate from Thailand (MW814863.1) possesses an insertion of 6 CAC repeats immediately upstream of the S9 stop codon (Figure S4A), resulting in a 122-codon gene. Almost all isolates also possess the S9-F3 AUG codon, initiating a 105-codon ORF (Figures S3, S4B). Two exceptions are the India|OM469318 isolate, which possesses a premature stop codon at position 50 and India|MG873154, which lacks the S9-F3 AUG. Thus, the S9-F3 gene is nearly ubiquitous amongst TiLV isolates and is also present in the divergent guppy TiLV-like virus.

As noted above, the moderate Kozak consensus surrounding the S9 AUG and the strong context of the S9-F3 AUG in the reference Israel Til-4-2011 (KU751822.1) strain we used here (Figure 1) suggests that S9-F3 may be expressed by leaky ribosomal scanning. To assess conservation of this potential translation mechanism, the Kozak consensus sequences flanking the S9 and S9-F3 start codons across all 47 sequences were analysed. Only 15 sequences included a sufficiently long 5′ UTR to permit full analysis of the Kozak consensus motif for the S9 AUG, and all possess an intermediate strength Kozak consensus (Figure 5B and S3). The remaining 32 sequences —though lacking complete Kozak data—consistently retained a +4 U nucleotide, indicating a generally moderate or weak Kozak context for S9. The uAUGs in the five Thai strains also displayed a +4 U (T), suggesting at best a moderate Kozak context, potentially affecting initiation at the downstream canonical S9. All 46 sequences with the S9-F3 AUG possess a strong Kozak context [-3 A, +4 G]. Collectively, these data support the hypothesis that S9-F3 is a conserved gene product likely expressed through leaky ribosomal scanning.

To investigate the evolutionary characteristics of the overlapping S9 and S9-F3 ORFs, we separately aligned 47 complete coding sequences for S9 and 45 for S9-F3 using Clustal Omega integrated in Jalview (Clamp et al., 2004) (Figure S4). The two sequences lacking an intact S9-F3 ORF (Figure S3) were excluded from the S9-F3 dataset. Residue-by-residue conservation scores (scored from 0 to 11), generated in Jalview based on biochemical property similarity (Livingstone & Barton, 1993), were extracted and normalised to a 0–1 scale for plotting. In the S9 protein, 69 out of 116 amino acids (59%) are fully conserved across all 47 strains (Figure 6A). The remaining positions exhibit varying degrees of variability, reflecting evolutionary divergence across the segment. For S9-F3, 69% of 105 residues are fully conserved, but unlike S9, where variable positions are scattered throughout the protein, conservation of S9-F3 is concentrated in the first 60 amino acids (Figure 6B). This more conserved region includes the predicted NES motif, although there is some variability within the NES itself. In contrast, the C-terminal region displays greater variability and is relatively rich in polar and charged residues. Consistent with this, it is predicted to be intrinsically disordered by the AIUPred programme (Erdős & Dosztányi, 2024), unlike the overlapping S9 protein (Figures 6C, D). To further assess evolutionary pressures, we employed the Fast Unconstrained Bayesian AppRoximation (FUBAR) method (Murrell et al., 2013) to the TiLV and TiLV-like sequences to estimate dN/dS ratios for each ORF, using the ratio of nonsynonymous [β] to synonymous [α] substitutions as a proxy. The β–α plot for S9 showed a clear signal of purifying (negative) selection, with ten codon positions (27, 45 67, 73, 81, 84, 90, 94, 98, 107) identified as statistically significant (Figure 6E, highlighted in red). This pattern supports the notion that S9 is a functionally essential and evolutionarily conserved protein. The S9-F3 β–α plot revealed five sites under significant negative selection, also scattered along the ORF (Figure 6F). Both genes also had three positions under positive (diversifying) selection and three of these (S9 codons 55 and 93 and S9-F3 codon 96) overlap sites of negative selection on the other gene. Thus, both S9 and S9-F3 are under evolutionary constraint, potentially reflecting important functions for both polypeptides.

**Figure 6.**
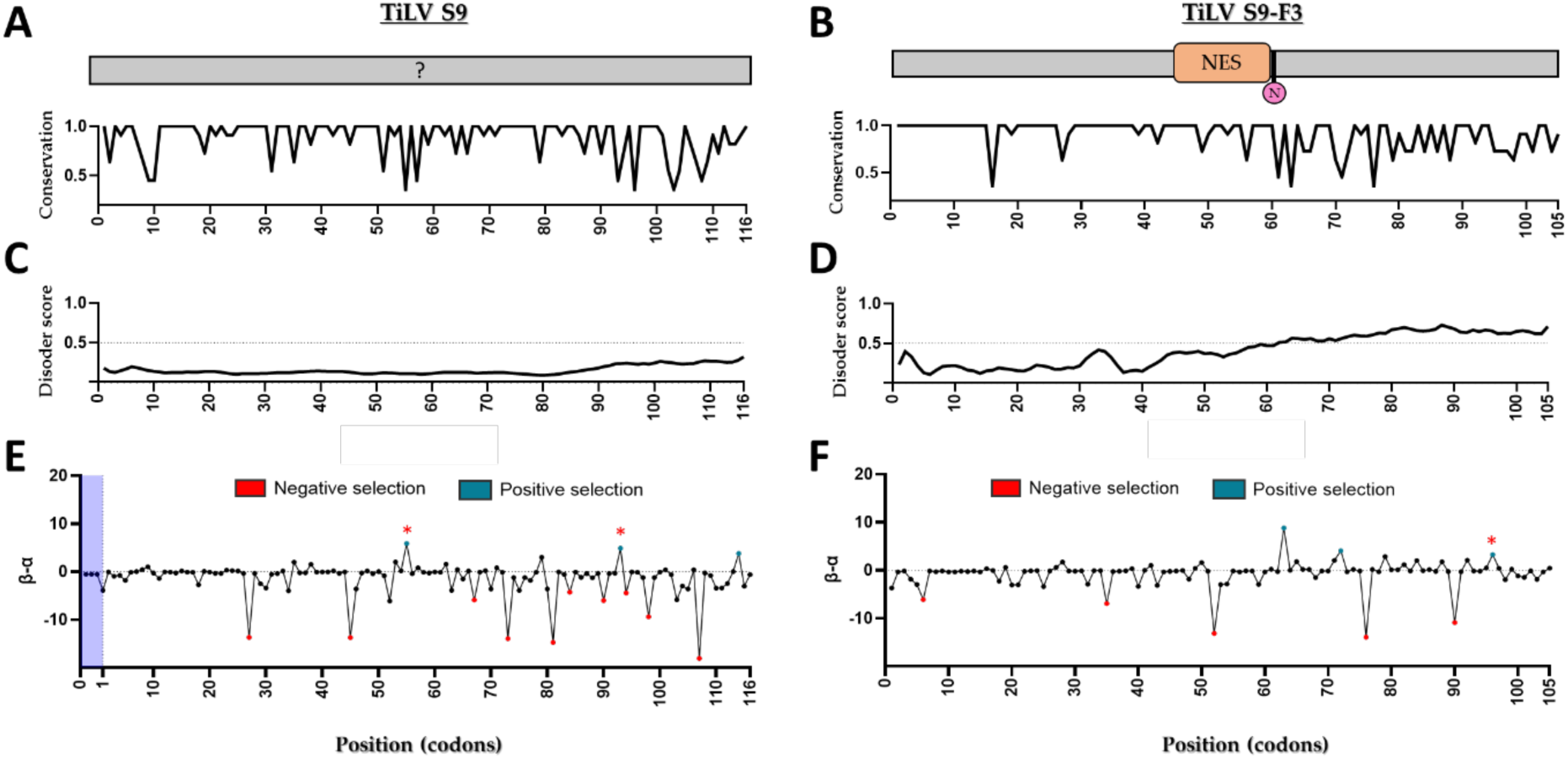
Sequence conservation of TiLV S9 and S9-F3 genes. (A, B) sequence conservation of TiLV S9 and S9-F3 proteins (1 = fully conserved). (C, D) Prediction of intrinsically disordered regions using the AIUPred algorithm. Values above 0.5 are suggestive of disorder. (E, F) Evolutionary selection pressure analysis of S9 and S9-F3 ORFs. The relative rate of nonsynonymous (β) to synonymous (α) substitutions was calculated using FUBAR (DataMonkey). Red points indicate significant purifying selection, blue points indicate diversification pressure. Red asterisks highlight where codons under significant pressure in the two ORFs overlap.

### Characterisation of major and alternative gene products of segment 9

Two distinct products were observed in cell-free translation reactions programmed with segment 9 (Figure 2): a discrete band (∼13 kDa) and another that ran more slowly as a broad smear, initially suspected to be an artifact caused by gel distortion resulting from the large amount of ß-globin in reticulocyte lysate (Pelham & Jackson, 1976). However, the discovery of S9-F3 in the MS data suggested that these two products might instead correspond to S9 and S9-F3 (with migration of the latter still affected by co-migration with unlabelled ß-globin). To test this, site-directed mutagenesis was used to create two mutants in which either the first AUG codon in frame 1 (ΔS9) or frame 3 (ΔS9-F3) was altered to UUG (leucine) (Figure 7A). The ΔS9-F3 mutation overlaps codon 3 of S9, but is silent. These constructs as well as WT segment 9 were then subjected to *in vitro* transcription and translation reactions followed by SDS-PAGE. Autoradiography showed that mutation of the canonical S9 AUG (ΔS9) resulted in a substantial decrease in the intensity of the 13 kDa band, while the smear product remained unaffected (Figure 7B). In contrast, mutation of the alternative S9-F3 AUG (ΔS9-F3) led to a substantial reduction in radiolabel intensity in the smear, and an increase in the smaller 13 kDa product. These findings suggest that the lower band (∼13 kDa) represents S9, whereas the slower-migrating smeared product corresponds to S9-F3. However, the slower migration of S9-F3 compared to S9, despite its lower predicted molecular weight, remains unexplained.

**Figure 7.**
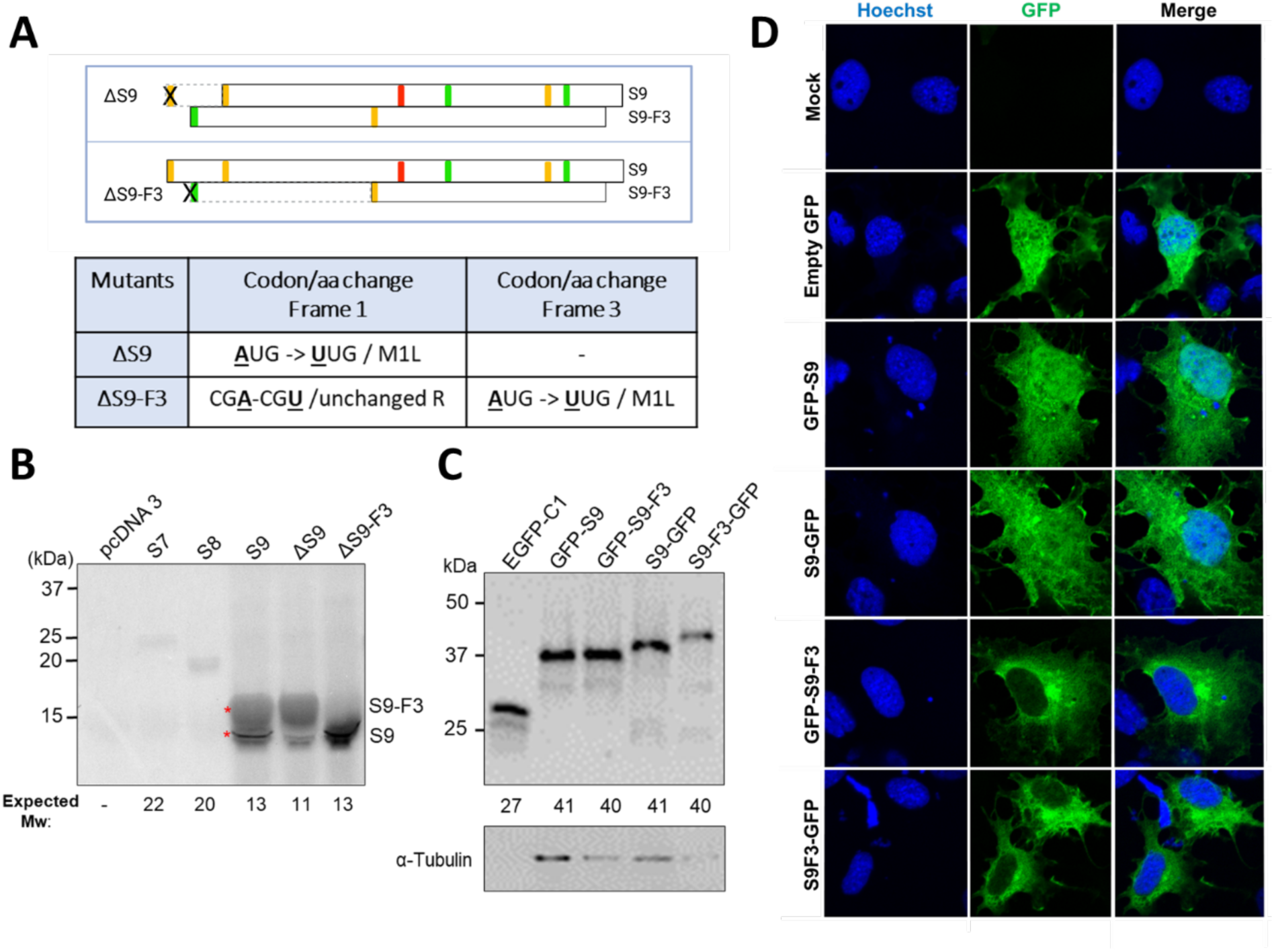
Molecular characterisation of primary and alternative gene products of segment 9. (A) Schematic representation of mutant segment 9 ORF structures, where the first AUGs from frame 1 or 3 were mutated (indicated by cross marks) to UUG (Leu) codons, with dashed boxes indicating portions of the ORFs now lacking a start codon. (B) *In vitro* translation of segment 9 proteins. Full-length coding sequences of empty vector, WT and mutated segments 7, 8 and 9 were transcribed and translated in rabbit reticulocyte lysate supplemented with [³⁵S]-methionine/cysteine. Translation products were resolved by SDS-PAGE and visualised by autoradiography. Asterisks indicate the two major products produced from segment 9. (C) Expression of GFP-tagged S9 and S9-F3 in cells. 293T cells were transfected with the indicated constructs, harvested 48 hours later and analysed by SDS-PAGE and western blotting for GFP or α-tubulin. (D) Intracellular localisation of WT S9 and S9-F3 tagged with GFP in tilapia cells. OmB cells transfected with the indicated plasmids were fixed 48 hours post-transfection, permeabilised and stained with Hoechst (blue) to delineate nuclei. Images (single optical slices) were acquired by a Zeiss 710 confocal microscope using a 63x objective and are representative of three independent experiments each performed with single technical repeat. Scale bars represent 10 μm.

To examine intracellular localisation of S9-F3, we created expression plasmids for N- and C-terminally tagged versions of S9-F3 and transfected them into 293T cells alongside the counterpart S9 plasmids. Both polypeptides expressed well with minimal levels of shorter than full-size products (Figure 7C), but similarly to the in vitro translation analysis, the GFP-tagged S9-F3 proteins either co-migrated in SDS-PAGE with the S9 protein (in the case of the N-terminally tagged versions) or migrated more slowly than the S9 fusion protein (as seen for the C-terminally tagged polypeptides). When transfected into OmB cells and imaged by confocal microscopy, the S9 and S9-F3 polypeptides showed distinct subcellular localisation patterns. Unlike the S9 protein, which as before, localised to both the nucleus and cytoplasm, the S9-F3 fusion proteins showed almost exclusively cytoplasmic localisation, with only minimal nuclear presence (Figure 7D). At around 40 kDa, monomeric S9-F3 polypeptides are substantially below the cut off size for proteins able to diffuse into the nucleus (Wang & Brattain, 2007), so this observation is consistent with the prediction of it containing an NES.

### Effects of a CRM1-dependent nuclear export inhibitor on S9-F3 localisation

To investigate whether nuclear export of the S9-F3 protein is mediated via the CRM1 (exportin 1) pathway, we used the CRM1-specific inhibitor Leptomycin B (LMB). LMB blocks CRM1-dependent nuclear export of various NES-containing proteins (Aumann et al., 2024; Fornerod et al., 1997) by covalently binding to a critical cysteine residue in CRM1, thereby preventing the formation of a stable export complex (Kudo et al., 1999; Kudo et al., 1998). To test if intracellular localisation of the TiLV polypeptides encoded by segment 9 were sensitive to CRM1 inhibition, 293T and tilapia OmB cells were transfected with plasmids encoding either S9 or S9-F3 fused to GFP, followed by treatment (or not) with LMB. Confocal microscopy revealed no significant change in the subcellular localisation of GFP alone or S9-GFP between LMB-treated and untreated cells, with signal distribution remaining both cytoplasmic and nuclear in either condition (Figure 8). In contrast, while S9-F3-GFP was primarily cytoplasmic in untreated cells of both species, LMB-treated cells displayed increased nuclear accumulation of S9-F3 in both tilapia and mammalian cell lines, particularly for the GFP-S9-F3 construct in 293T cells. These findings indicate that the S9-F3 protein undergoes active nuclear export mediated via the CRM1 pathway, consistent with the presence of a predicted NES within its sequence.

**Figure 8.**
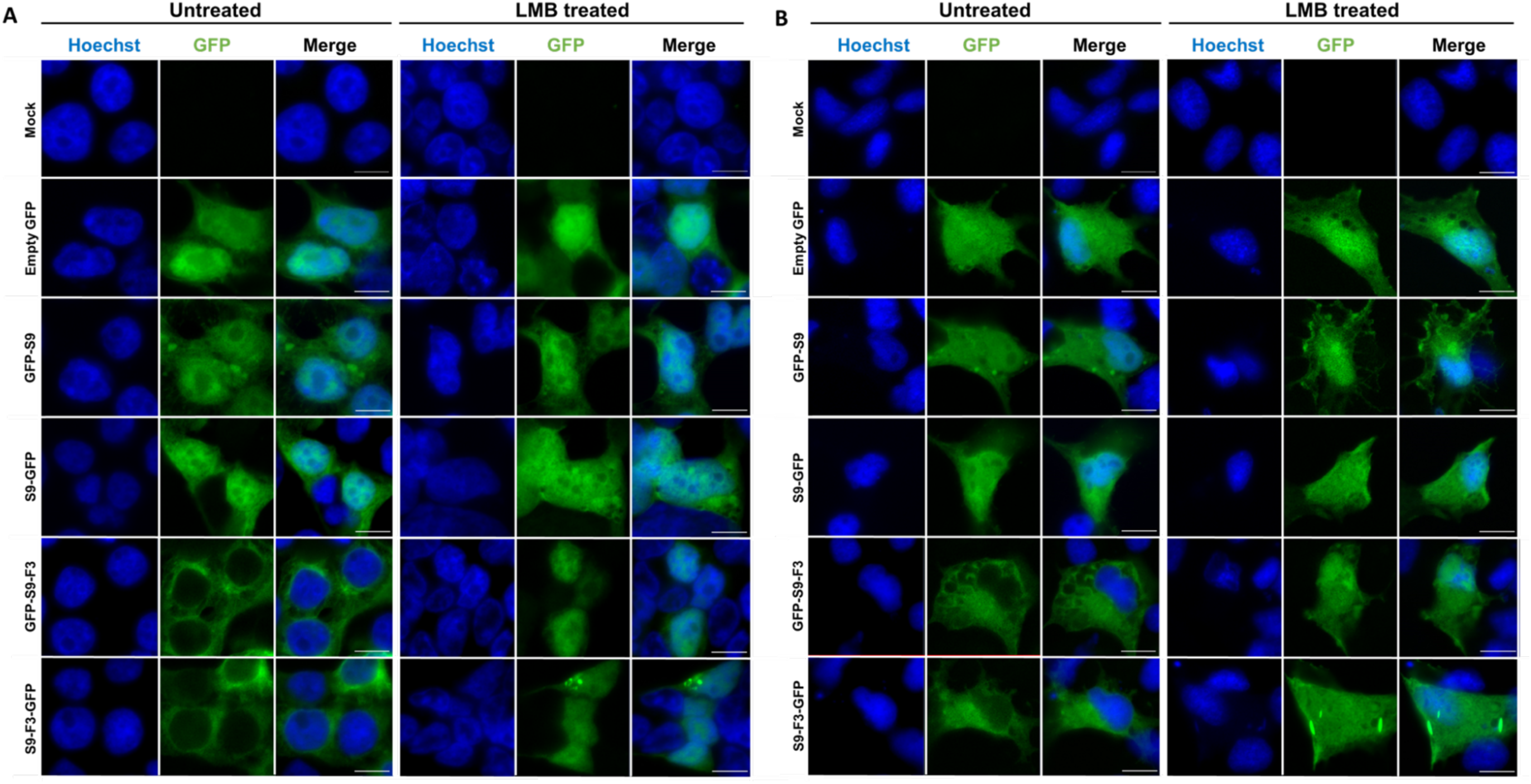
Effect of LMB on nuclear export of the TiLV S9-F3 protein. Mammalian 293T (A) and tilapia OmB (B) cells were transfected with the indicated plasmids (or mock-transfected) for 24 hours prior to LMB treatment (10 nM). Following 24 hours of LMB exposure, cells were fixed and nuclei were stained with Hoechst dye (blue). Image acquisition was carried out on a Zeiss 710 confocal microscope employing a 63x objective. Images are single optical slices representative of three independent experiments, each conducted with a single technical repeat. Scale bars equal 10 μm

## Discussion

This study provides the first expression and subcellular localisation analysis of the TiLV proteome, encompassing the primary products of all ten genomic segments. By combining *in vitro* translation and cell-based expression systems with proteomic validation from virus-infected cells, we confirmed the expression of all ten canonical TiLV proteins and discovered a previously unreported product, S9-F3, translated from the second reading frame of segment 9.

Using rabbit reticulocyte lysate-based *in vitro* translation of transcripts encoding untagged, wild-type TiLV proteins, as well as transfecting cells with plasmids encoding GFP-tagged version, we successfully expressed proteins of approximately the expected sizes from the plus strand of most TiLV segments. Segment 10 was an exception, consistently producing a polypeptide that exhibited anomalously slow migration on SDS-PAGE in both cell-free and GFP-tagged cellular expression systems. This unusual migration is consistent with findings by Yang et al. (2024), who reported similar behaviour after expressing S10 in *Escherichia coli*. The highly acidic isoelectric point of the S10 protein likely contributes to this effect, as small, acidic proteins often bind SDS poorly, resulting in the inaccurately apparent molecular weights (Nelson, 1971; Nowakowski et al., 2014). The second polypeptide encoded by segment 9, S9-F3, also migrated more slowly in SDS-PAGE gels than expected, although it has a slightly basic predicted pI of 7.8. Whether its aberrant migration results from some form of post translational modification remains to be determined. The protein encoded by segment 5 also exhibited some peculiarities in SDS-PAGE; it was readily expressed *in vitro* and migrated as expected in gels, but when expressed in cells as a GFP-tagged fusion protein, it was poorly detectable by western blot. Further analysis to be reported elsewhere (Pankaew et al., manuscript in preparation) suggests this may be due to heat-induced aggregation of the polypeptide during preparation for SDS-PAGE exacerbated by its entry into the cellular secretory pathway, which does not occur in reticulocyte lysate.

Additional polypeptides detected in *in vitro* translation reactions from segments 1, 5, 6, and 9 suggest potential alternative translation initiation, perhaps arising via leaky ribosomal scanning. In segment 1, a weak Kozak context around the canonical start codon followed by a tryptophan codon may lead to reduced expression and stability of the primary product (Varshavsky, 1996), favouring the accumulation of alternative isoforms initiated from stronger downstream AUGs. This pattern of a primary methionine codon followed by a destabilising residue also occurs in segments 5 and 6, though is further complicated by the possible presence of signal peptide sequences. In the case of S1, two peptides likely to arise from initiation at the first methionine codon were detected by MS, while for S5 and S6 only peptides downstream of the second or third AUG codon respectively were detected. Exact definition of what polypeptide species are produced from these segments therefore requires further analysis.

A computational prediction has suggested up to 14 potentially functional ORFs from both polarities of the TiLV genome (Acharya et al., 2019), but without experimental validation. Mass spectrometry of virus polypeptides from infected OmB cells confirmed only one additional low-abundance protein—S9-F3—beyond the ten canonical proteins. While Acharya et al. (2019) proposed a related ORF (S9ORF2), their analysis did not require initiation at an AUG codon. In contrast, our *in vitro* translation and segment mutagenesis data suggest that S9-F3 originates from a methionine-initiated ORF from reading frame 3 embedded entirely within the S9 gene, probably expressed via leaky ribosomal scanning. Although our proteomics detected only one additional product, this does not exclude the possibility of other low-abundance polypeptides being expressed, warranting further investigation.

Analysis of all available S9 sequences in public databases revealed strong conservation of the S9-F3 ORF across isolates, extending to a TiLV-related virus from guppy fish. Multiple sequence alignment of S9-F3 revealed strong conservation, particularly within the N-terminal region (amino acids 1–60), where a putative nuclear export signal (NES) is located, supporting its potential as a functional (although perhaps not essential, given two viruses with a disrupted S9-F3 ORF) component of the TiLV life cycle. The potential NES within S9-F3 and its exclusively cytoplasmic localisation pattern led us to investigate whether it undergoes active nuclear export. Leptomycin B treatment indicates that S9-F3 does indeed utilise CRM1-dependent nuclear export. This is reminiscent of nuclear export proteins (NEPs) in orthomyxoviruses such as IAV (Boulo et al., 2007; Huang et al., 2013) and infectious salmon anaemia virus (Ritchie et al., 2002; Zhang et al., 2017), which express NEPs from spliced mRNAs of segment 8 and segment 7, respectively. In IAV, NEP consists of a flexible N-terminal domain (aa 1–53) and a structured C-terminal domain (aa 54–121), with CRM1-dependent NES motifs located in the N-terminal unstructured region (Huang et al., 2013; Iwatsuki-Horimoto et al., 2004; O’Neill et al., 1998). No structural information is available for TiLV S9-F3 but it may also consist of structured and unstructured domains (albeit the other way round to NEPs of orthomyxoviruses), with the NES motif located near the domain boundaries. It also differs in being generated (probably) via leaky scanning of an unspliced transcript, in a gene organisation reminiscent of ISAV segment 8, which encodes a matrix protein and an interferon antagonist (García-Rosado et al., 2008). We also note that the C-terminus of S9-F3 contains a PDZ-binding motif, similarly to the NS1 interferon antagonist of some strains of IAV (Jackson et al., 2008; Lee & Zheng, 2010). Future work is needed to determine the function of S9-F3 and whether it is essential for TiLV replication or functions as an accessory protein.

In IAV, nuclear import of the viral RNA synthesis machinery relies on NLS motifs in the polymerase subunits and NP. PB2 and NP independently localise to the nucleus (Mukaigawa & Nayak, 1991; Wang et al., 1997), while PB1 and PA require cytoplasmic dimerisation prior to nuclear import (Fodor & Smith, 2004). In TiLV, *in situ* hybridisation studies identified the nucleus as the primary site of transcription (Bacharach et al., 2016), suggesting a similar need to import S1-3 polypeptides into the nucleus. Our localisation data for the individually expressed RNP subunits lead to speculation of a similar mechanism for import of the TiLV P proteins. S2 (PB2-like) was predominantly nuclear and contains a potential C-terminal NLS, consistent with IAV PB2. In contrast, S1 (PB1-like) and S3 (PA-like) were primarily cytoplasmic, consistent with the behaviour of their IAV counterparts before dimerization. S4 (NP-like) displayed both nuclear and cytoplasmic localisation, aligning with previous reports on exogenously expressed IAV NP (Digard et al., 1999) and recent findings by Abu Rass et al. (2022).

In conclusion, our findings confirm expression of all canonical TiLV proteins, identify a novel NEP-like protein with active nuclear export features, and provide the first TiLV proteome-wide localisation map. These insights not only enhance understanding of TiLV molecular biology but also hint at evolutionarily conserved mechanisms among segmented negative-sense RNA viruses in aquatic hosts. The discovery of S9-F3 underscores the potential for alternative translation strategies to drive functional diversification and offers a foundation for future investigations into TiLV pathogenesis and antiviral strategies.

## Materials and Methods

### Computational biology analyses of TiLV segments

Sequence information for the ten TiLV segments from strain Til-4-2011 (assession nos. KU751814.1-KU751823.1) was retrieved from the National Center for Biotechnology Information (NCBI) database and analysed for their nucleotide (nt) and untranslated region (UTR) length using Benchling software (Benchling, 2020). ORF finder (Rombel et al., 2002) was employed to identify the primary and alternative plus strand ORFs in each segment, initiated with an AUG codon and a minimum length of 25 codons. The Kozak consensus sequence of these ORFs was assessed based on the presence or absence of a purine (A/G) at position -3 and a G at position +4, with the A of the AUG codon designated as +1 (Kozak, 1986; Meijer & Thomas, 2002). If both positions matched consensus, the AUG was classified as a strong context, if neither, weak and if one or other of +1 or -3 matched, as intermediate (Wise et al., 2009). Potential functional motifs in the primary ORFs, including nuclear localisation signal (NLS), signal peptide (SP), SP cleavage site (SPC), N-linked glycosylation (N-gly), transmembrane domain (TMD), and nuclear export signals (NES were predicted using cNLS Mapper (Kosugi et al., 2008), SignalP 5.0 (Almagro Armenteros et al., 2019), NetNGlyc - 1.0 (Gupta & Brunak, 2002), TMHMM - 2.0 (Krogh et al., 2001; Sonnhammer et al., 1998) and LocNES (Xu et al., 2015). The prediction confidence was assessed based on output probability scores generated by each software tool. For NLS prediction, a cutoff score of 5 was applied (scoring range: 1–10, where 1–2 = cytoplasmic, 3–5 = nuclear & cytoplasmic, 6–7 = partially nuclear, 8–10 = predominantly nuclear). For SP, N-gly, and TMD, a cutoff score of 0.5 was used, while for NES, a cutoff of 0.3 (on a 0–1 scale) was applied, with higher scores indicating higher confidence in prediction accuracy.

### Cell culture

Human Embryonic Kidney 293T (293T) cells (DuBridge et al., 1987) were maintained in Dulbecco’s modified Eagle medium (DMEM), supplemented with 10% foetal bovine serum (FBS), 100 U/ml penicillin and 100 µg/ml streptomycin at 37°C and 5% CO_2_. Immortalised cells (OmB) from the brain of tilapia *(Oreochromis mossambicus)* (Gardell et al., 2014) were cultured in L-15 medium (Sigma) with 10% FBS, 100 U/ml penicillin and 100 µg/ml streptomycin at 28°C without controlled CO_2_. Regular cell passage was performed when the cells reached 80% confluence.

### Plasmids

Synthetic TiLV cDNAs, corresponding to genome segments 1–10 (GenBank accession numbers KU751814–KU751823), were synthesised by Integrated DNA Technologies (IDT) and cloned into the pMiniT 2.0 vector (New England Biolabs) at the toxic Minigene cloning site downstream of a bacteriophage T7 RNA polymerase promoter for *in vitro* protein expression. To generate GFP-tagged constructs, viral cDNA segments were subcloned into pEGFP-C1 and pEGFP-N3 expression vectors (Invitrogen) using the Gibson Assembly method (Gibson et al., 2009). Site-directed mutants were created using the QuikChange II Site-Directed Mutagenesis Kit (Agilent Technologies), following the manufacturer’s instructions. All oligonucleotides used for cloning and mutagenesis are listed in Supplementary Tables S6 and S7. The integrity and accuracy of all constructs were confirmed by Sanger sequencing (Genewiz).

### Virus propagation and sample preparation for proteomic analysis

TiLV strain TH-2018-N was kindly provided by Prof. Ha Thanh Dong (Asian Institute of Technology, Thailand). Viral stocks were propagated by infecting confluent monolayers of OmB cells at a MOI of 0.01. Infected cultures were incubated for up to 10 days at 25 °C or until clear cytopathic effects (CPE) were observed. Supernatants were harvested, clarified by centrifugation at 1,500 rpm for 5 minutes to remove cellular debris, aliquoted, and stored at −80 °C for subsequent titration via TCID₅₀ assay. Cell pellets collected at 10 days post-infection (dpi) were resuspended in 50 μL of phosphate-buffered saline (PBS) and submitted to the Roslin Institute Proteomics and Metabolomics Facility for proteomic analysis using liquid chromatography–mass spectrometry (LC–MS).

### In vitro transcription and translation (IVTT) of TiLV proteins

Coupled *in vitro* transcription and translation (IVTT) reactions were performed in rabbit reticulocyte lysate using the TnT® T7 Quick Coupled Transcription/Translation System (Promega). Each reaction (10 µL) contained 200 ng of plasmid DNA encoding individual TiLV gene segments or an empty vector control, supplemented with EXPRESS ^35^S3 Protein Labeling Mix of [³⁵S]-L-methionine and [³⁵S]-L-cysteine (PerkinElmer) to enable radiolabelled protein synthesis. Reactions were carried out according to the manufacturer’s protocol. Following translation, radiolabelled proteins were mixed with 2×Laemmli sample buffer, denatured at 95 °C for 10 minutes, and resolved on 15% SDS–polyacrylamide gels. Gels were fixed in a methanol/acetic acid fixing solution for three 10-minute intervals under constant agitation, then dried under vacuum at 80 °C for 2 hours. Dried gels were exposed to X-ray film in a light-tight cassette overnight and developed using a Konica SRX-101A X-ray film processor (FAPJ).

### SDS-PAGE and western blot analysis

Cells transfected with individual GFP-tagged TiLV expression plasmids were harvested in Laemmli sample buffer (Bio-Rad). Lysates were denatured by heating at 95 °C for 10 minutes. Proteins were resolved by SDS–PAGE using precast 4–15% gradient gels (Bio-Rad) and transferred onto nitrocellulose membranes (Invitrogen) using the Trans-Blot Turbo™ Transfer System (Bio-Rad). Membranes were blocked with 5% non-fat dry milk in PBS containing 0.1% Tween-20 (PBS-T) for 1 hour at room temperature, followed by incubation with primary antibodies—anti-GFP (clone JL-8; Takara, 632381) and anti-α-tubulin (LifeTechnologies, MA180017) diluted in blocking buffer overnight at 4 °C. After three 5-minute washes with PBS-T, membranes were incubated with species-specific IRDye-conjugated secondary antibodies (LI-COR Biosciences) for 1 hour at room temperature in the dark. Following additional PBS-T washes, signals were detected and imaged using the LI-COR Odyssey imaging system.

### Mass spectrometry analysis

Protein samples were prepared using S-Trap™ micro spin columns (Protifi, Fairport, USA) following the manufacturer’s protocol. Protein concentrations were determined using the bicinchoninic acid (BCA) assay (ThermoFisher Scientific). Samples were reduced with 5 mM dithiothreitol (DTT) and alkylated with 20 mM iodoacetamide (IAA). Phosphoric acid was then added to achieve a final concentration of 2.5%, followed by protein clean-up via desalting in wash buffer. Proteins were digested with trypsin, and the resulting peptides were enriched and desalted prior to elution using a buffer designed to recover both hydrophilic and hydrophobic peptides. Purified peptides were separated using an 85-minute linear gradient on an Aurora 25 cm analytical column (IonOpticks, Australia) with an UltiMate RSLCnano LC system (ThermoFisher Scientific), coupled to a timsTOF fleX mass spectrometer via a CaptiveSpray ionization source. The flow rate during separation was maintained at 200 nL/min and increased to 500 nL/min during column washout. The column temperature was held at 50 °C. Data-dependent acquisition with parallel accumulation–serial fragmentation (DDA-PASEF) was used. Full MS scans were acquired from 100–1700 m/z across a 1/K₀ ion mobility range of 1.45 to 0.65 Vs/cm². Up to 10 PASEF MS/MS scans were performed per acquisition cycle, targeting precursors with charges >1 and applying an intensity threshold of 1,750 counts and a target intensity of 14,500 counts. Raw MS data were analysed using PEAKS Studio 11.5 (Bioinformatics Solutions Inc., Ontario, Canada), searching against the *Oreochromis niloticus* reference proteome (RefSeq: GCF_001858045.2) and a custom six-frame translation of the TiLV genome including open reading frames ≥9 amino acids. Peptides were identified with standard modifications and one missed cleavage, and protein abundance was estimated using label-free quantification based on extracted ion chromatograms. Extracted ion chromatograms (XICs) of peptide precursors were integrated over time to determine the area under the curve (AUC), which was used to estimate relative protein abundance.

### Fluorescence imaging of protein localisation

293T or OmB cells were seeded on fibronectin-coated 10 mm glass coverslips in a 24-well plate and transfected with individual plasmid constructs using Lipofectamine 2000 (Thermo Fisher Scientific), according to the manufacturer’s instructions. At 48 hours post-transfection, cells were fixed with 4% paraformaldehyde for 20 minutes at room temperature, followed by three washes with phosphate-buffered saline (PBS). Permeabilisation was performed using PBS containing 0.2% Triton X-100 for 10 minutes at room temperature, followed by an additional three PBS washes. Nuclei were stained with Hoechst 33342 (1:5000 dilution in blocking buffer) for 10 minutes in the dark at room temperature. After staining, coverslips were washed three times with PBS and mounted onto glass slides using ProLong™ Gold Antifade Mountant (Thermo Fisher Scientific). Confocal fluorescence imaging was performed using a Zeiss LSM 710 laser scanning confocal microscope equipped with a 63× oil immersion objective. During acquisition, identical imaging settings were maintained across all experimental conditions, including laser power, detector gain, and scan parameters for each fluorophore. Multi-channel acquisition and microscope control were conducted using ZEN 2012 (Black edition) software. Image processing, including scale bar addition and fluorescence channel visualisation, was performed using Fiji (ImageJ). All experiments were independently repeated at least three times.

### Nuclear export inhibition using Leptomycin B (LMB)

To investigate nuclear export inhibition, LMB was used in a CRM1 inhibition assay. OmB and 293T cells were transfected with S9 and S9-F3 expression constructs (and controls) for 24 hours, followed by incubation with either 10 nM LMB or dimethyl sulfoxide (DMSO; vehicle control) for another 24 hours. After 48 hours of transfection, cells were processed for confocal microscopy as described above.

### Conservation analysis of S9-F3

A total of 53 TiLV and TiLV-like segment 9 genome (in plus sense cDNA format) and protein sequences, originating from 12 countries, were retrieved from the NCBI TiLV sequence database as of May 2025. ORFs were identified using the NCBI ORF Finder tool to extract putative S9-F3 protein sequences. To ensure inclusion of only full-length coding sequences for downstream analyses, initial alignments were performed using the Clustal Omega web service within Jalview (Clamp et al., 2004) under default parameters. Of the 53 sequences, 6 were excluded due to incomplete 5′-end cDNA coverage (missing 20–44 nucleotides): PP600914.1, KY817383.1, KY817382.1, KY817381.1, MN061920.1, and MW591475.1. The remaining 47 complete sequences were retained for MSA and conservation analysis of segment 9 (Table S5). In addition, the sequence for Thailand|MW814863 includes an unusual repeat of 6 x CAC codons just before the S9 stop codon. For S9-F3-specific analysis, an equivalent approach was applied. Seven sequences were excluded: the six incomplete sequences mentioned above and one additional sequence (MG873154.1) in which the first AUG codon was substituted by CCG, preventing predicted S9-F3 translation. Another sequence (OM469318.1) was excluded due to the presence of a premature stop codon at amino acid position 50. Consequently, 45 complete S9-F3 protein sequences were used for conservation analysis. Following MSA, three annotations were generated: sequence consensus, alignment quality, and conservation scores. Conservation was numerically scored on a scale where a score of 11 indicated fully conserved residues; a score of 10 represented mutations with conserved biochemical properties; and scores below 10 indicated lower conservation. These scores were normalised to a 0–1 scale and visualised as a conservation plot (X-axis: amino acid position; Y-axis: normalised conservation score) using GraphPad Prism.

### Phylodynamics and temporal phylogenetics analyses

Multiple phylogenetic and phylodynamic models were applied to map the evolution and spread history of TiLV. Preliminary phylogenetic trees were generated using a maximum likelihood model using IQTree and the temporal signals of the sequence data were assessed using TempEst v1.5.3. The MW814863 sequence with 6 repeats of CAC codons just before the S9 stop codon was edited out for phytogenic analyses. A time-scaled phylogeny of TiLV was produced using Bayesian phylogenetic methods in BEAST version 1.10.4 (Suchard et al., 2018) with the TN93+G substitution model, a strict molecular clock, and an exponential population growth prior, based on 150 million posterior trees. Metadata on geography and host was collected from NCBI. Additionally, the presence or absence of a uAUG in TiLV S9 was annotated post-alignment and used to perform phylogenetic and phylogeographic analyses. The resulting annotated, time-scaled phylogenetic tree was visualised in R, using the ggtree package (Xu et al., 2021).

## Data availability

The data supporting this study are available from the corresponding authors upon reasonable request. The mass spectrometry proteomics data have been deposited to the ProteomeXchange Consortium via the MASSIVE partner repository with the dataset identifier PXD68424 and will be made public on peer reviewed publication.

## Author Contributions

N.P. conceptualisation, methodology, investigation, data curation, validation, writing – original draft preparation, review & editing, visualization, funding acquisition;

D.K. investigation, data curation, writing – review & editing;

F.A. data curation, writing – review & editing;

J.P. supervision, writing – review & editing;

R.H. resources, supervision;

R.G. resources, supervision, writing – review & editing;

R.M.P. methodology, data curation, writing – review & editing;

P.D. conceptualisation, writing—review and editing, supervision, project administration funding acquisition.

## Acknowledgements

We would like to thank Profs. Kim Thompson and David Bauer for helpful advice and discussion. We thank Dr Dietmar Kultz (University of California, Davis) and Asst Prof. Ha Thanh Dong (Asian Institute of Technology, Thailand) for providing OmB cells and TiLV strain TH-2018-N, respectively. We also acknowledge the Roslin Bioimaging staff for their continued guidance and timely support with imaging analysis and Judit Aguilar for help with the mass spectrometry. N.P. gratefully acknowledges Kamphaeng Phet Rajabhat University, Thailand, for supporting her academic development through the Royal Thai Government Scholarship scheme.

## Supplementary figures and Tables

**Figure S1.**
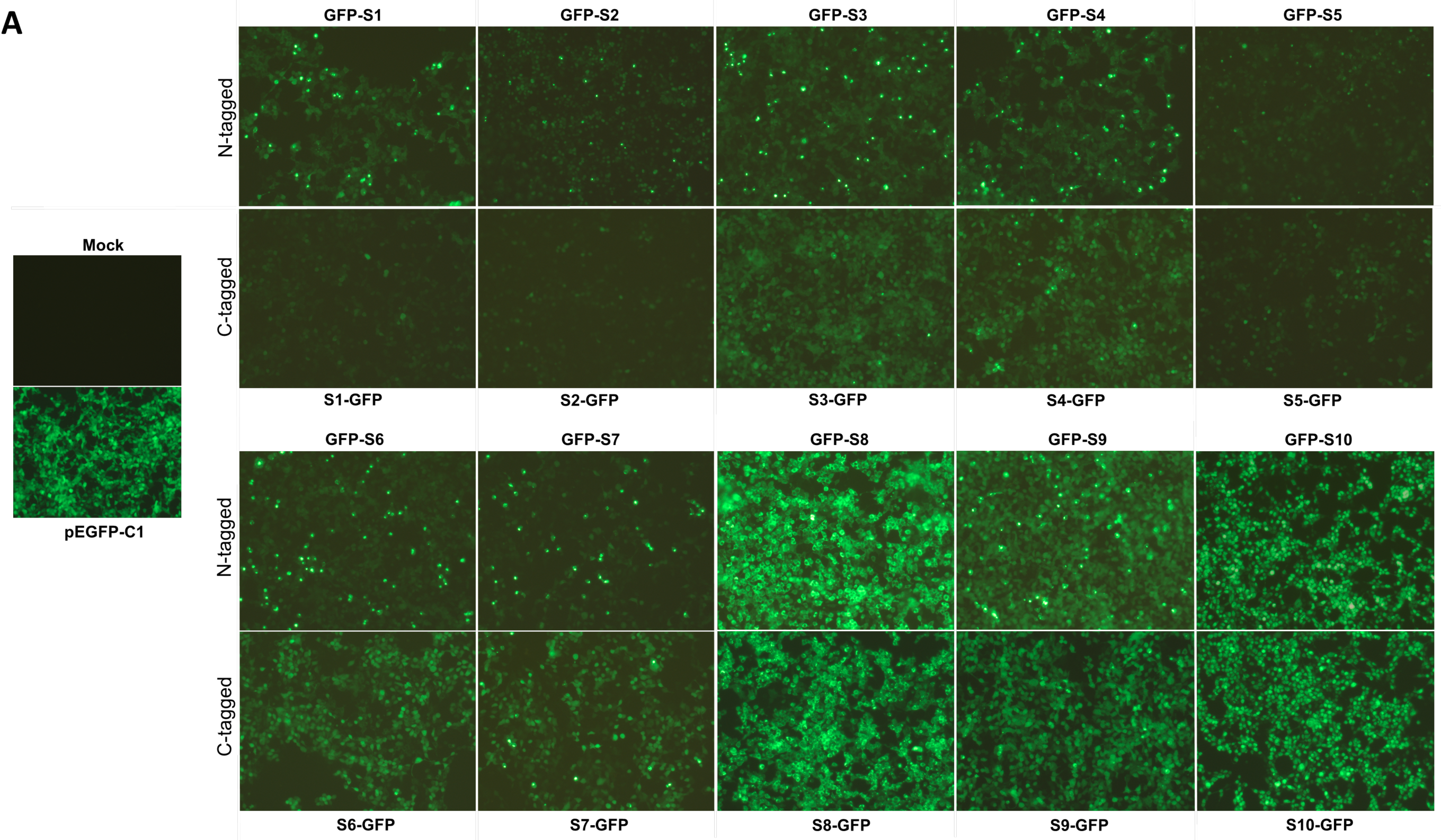
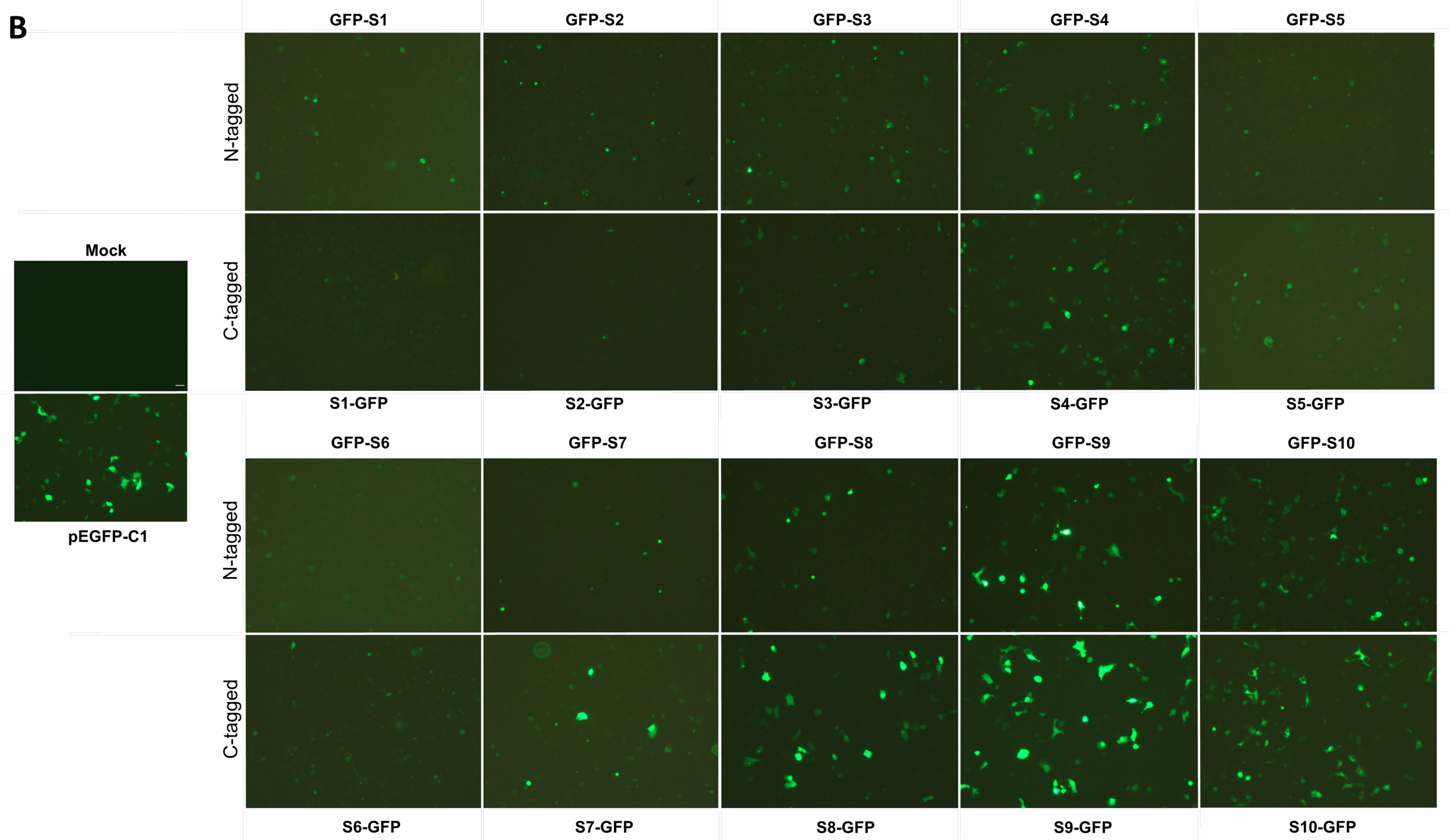
Transfection efficiency of GFP-fusion S1-S10 proteins in mammalian and tilapia cells. 293T (A) and OmB (B) cells were transfected with indicated plasmids expressing TiLV proteins or empty GFP plasmid or mock controls. After 48 hours of transfection, live cells were visualised for positive GFP signal under a fluorescent microscope (Zeiss Axiovert 25) and images were acquired with a 10x objective.

**Figure S2.**
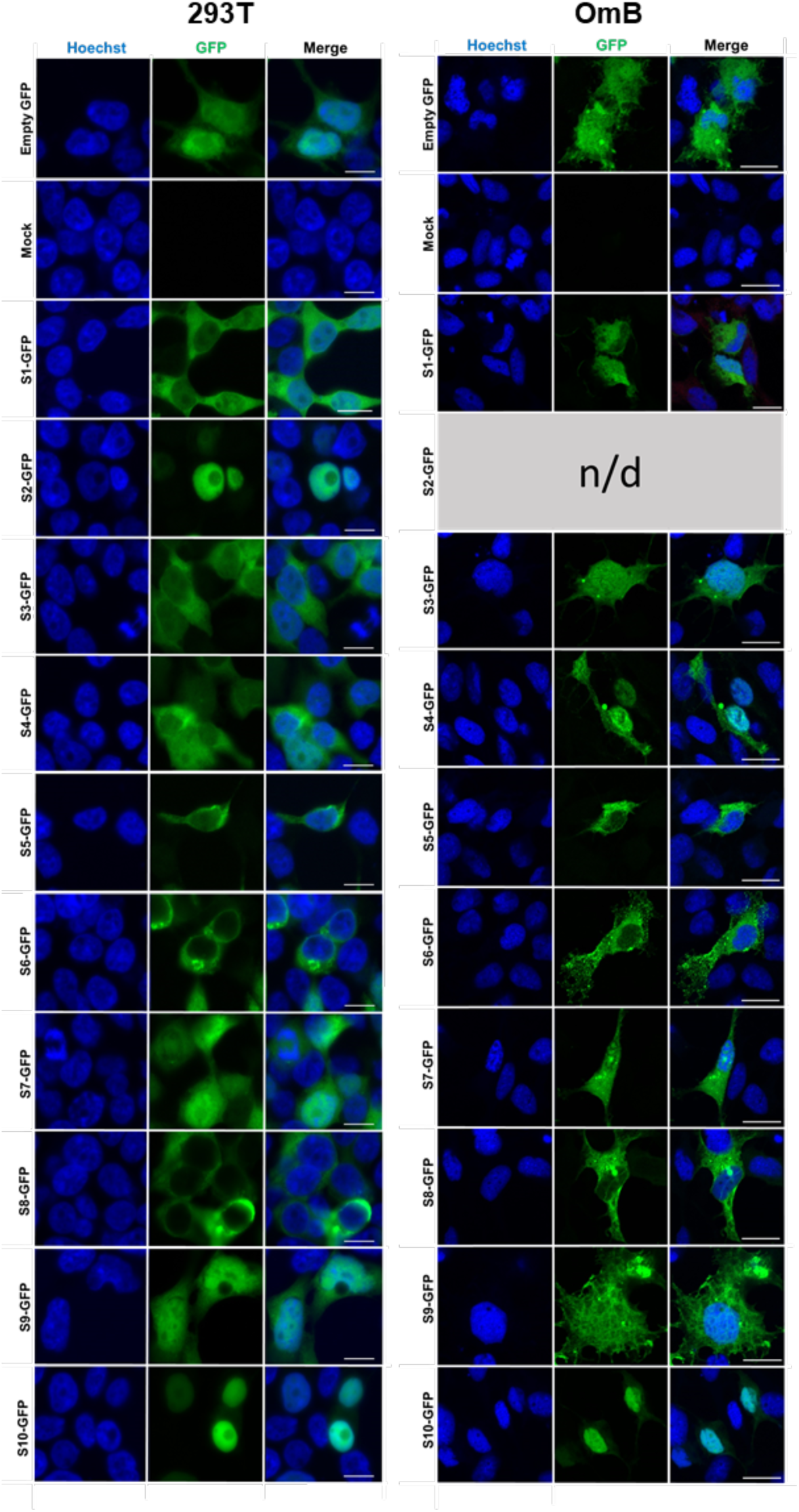
Subcellular localisation of GFP-tagged TiLV proteins in mammalian and tilapia cells. (A) 293T and (B) OmB cells were transfected with plasmids expressing C-terminal GFP fusion constructs of TiLV segments 1–10 or an empty GFP vector (pEGFP-C1), or were mock-transfected. 48 h post-transfection, cells were fixed and stained with Hoechst dye to visualise nuclei. GFP (green) and Hoechst (blue) fluorescence images were acquired using a Zeiss LSM710 microscope. Images are single optical slices, representative of three independent experiments, each performed with a single technical replicate. Scale bars: 10 µm, n/d: not determined because of poor expression.

**Figure S3.**
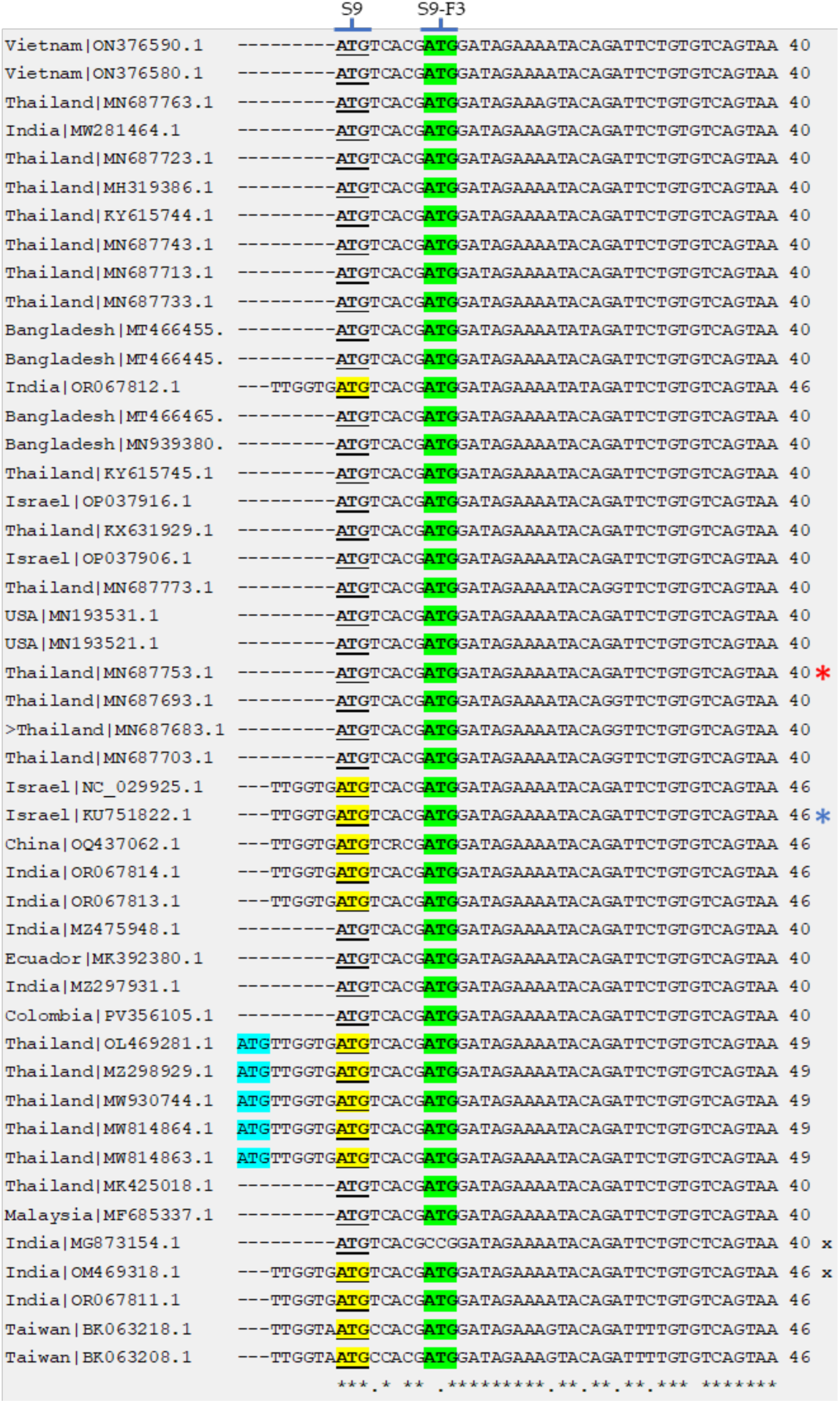
Conservation analysis of TiLV S9 5′ sequences and Kozak consensus motifs. Forty-seven segment 9 sequences (plus sense, given as cDNA) were retrieved from the NCBI database and aligned using Clustal Omega, with visualization and annotation performed in Jalview. A partial alignment of the 5′ region is shown, with residue positions indicated at the end of each sequence. Conservation at each nucleotide position is marked by black asterisks below the alignment. The canonical start codons (ATG) for the primary S9 ORF and the alternative S9-F3 ORF are highlighted, with yellow and green indicating intermediate and strong Kozak consensus strength, respectively. Additional upstream ATG codons (highlighted in blue) represent in-frame alternative start sites with, at most, intermediate Kozak consensus. Cross symbols denote two sequences excluded from the S9-F3 alignment due to the presence of a premature stop codon or the absence of an AUG initiation codon. Red and blue asterisks indicate sequences corresponding to the virus isolate and synthetic clones used in this study, respectively.

**Figure S4.**
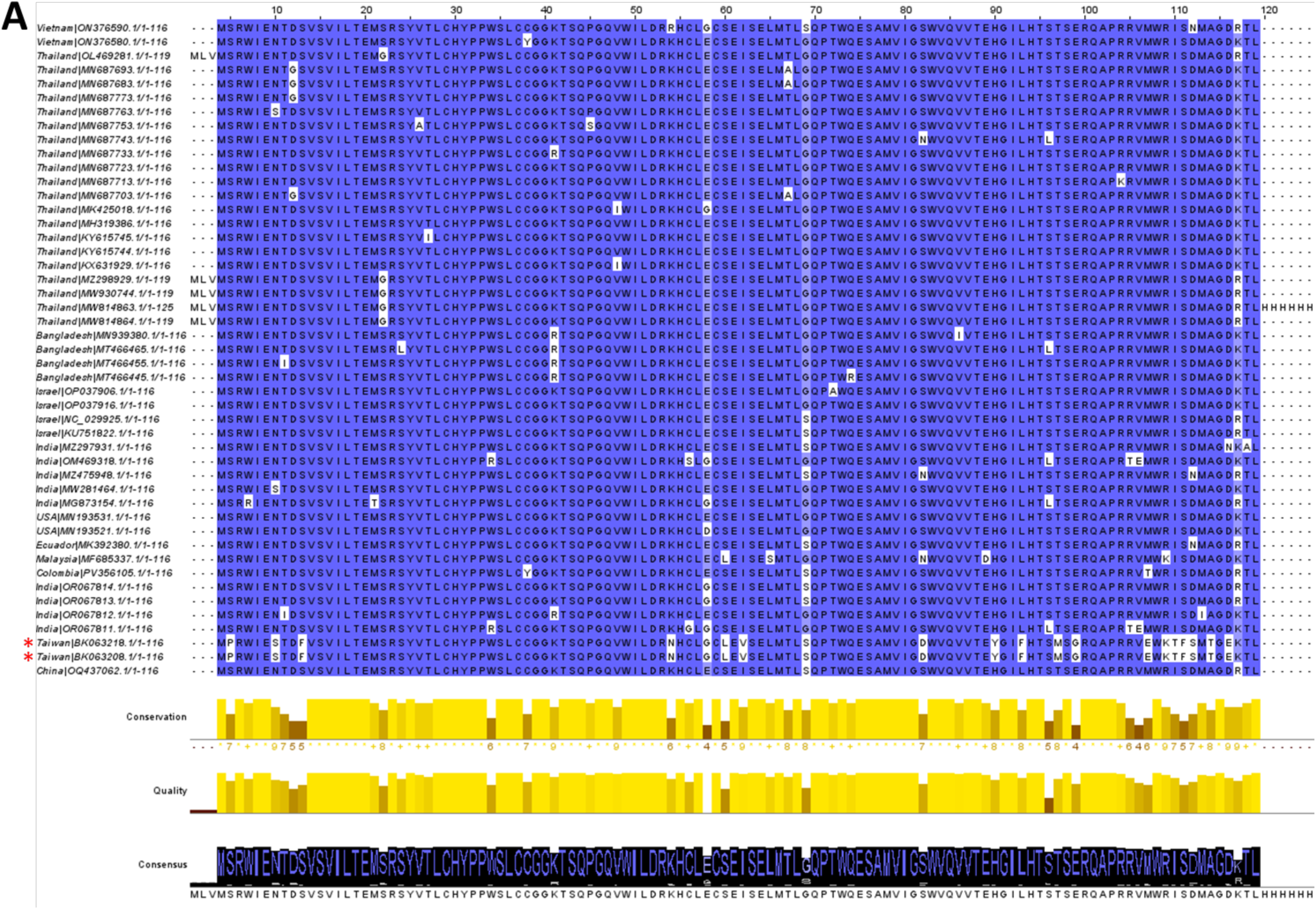
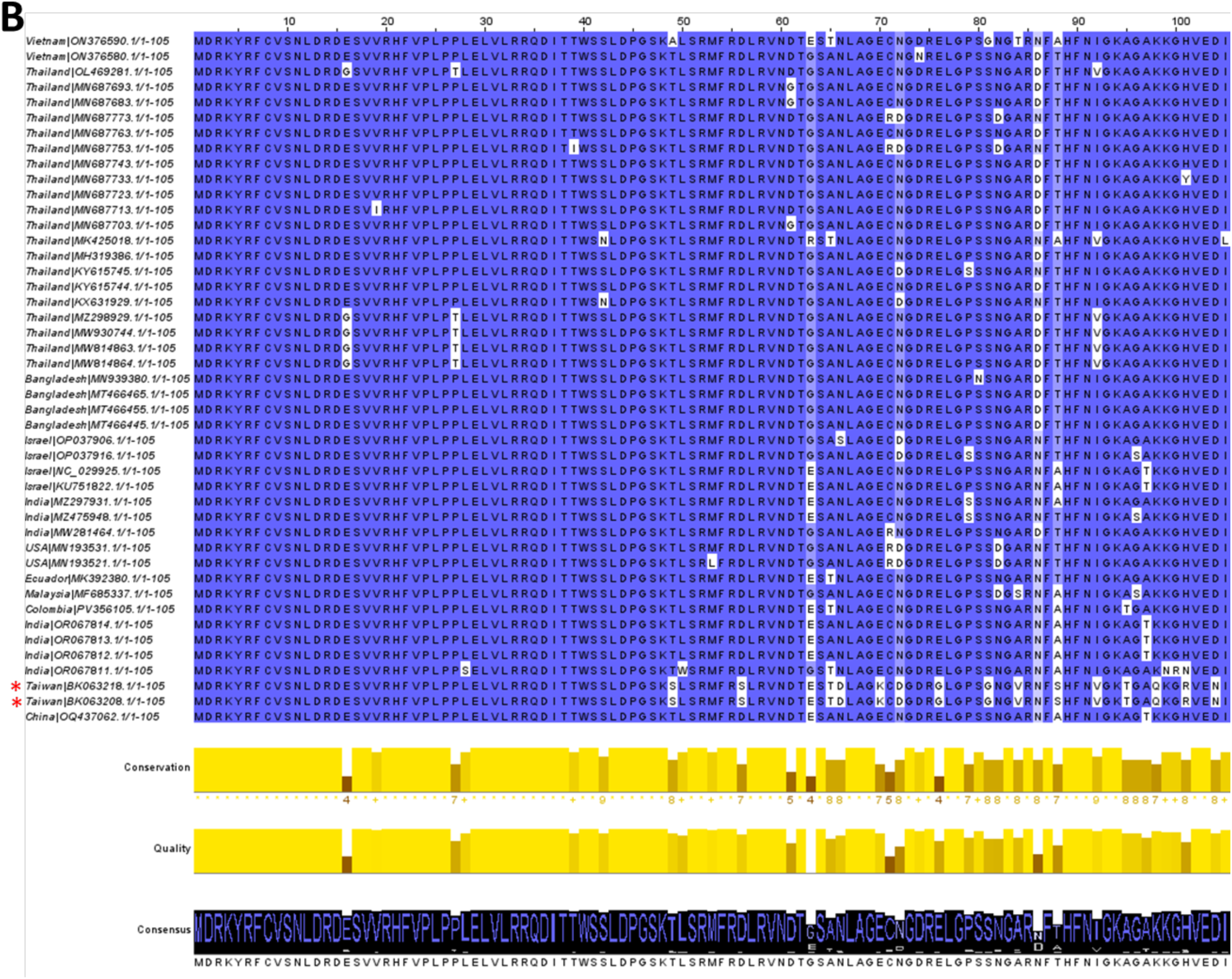
Multiple sequence alignment of TiLV S9 and S9-F3 predicted polypeptide sequences. 47 complete coding sequences of S9 (A) and 45 of S9-F3 (B) were retrieved from the NCBI database and aligned via Clustal using Jalview software. Red asterisks indicate the guppy TiLV-like isolates from Taiwan. Aligned residues are color-coded based on conservation; a stronger intensity indicates a higher physicochemical property conservation score. Consensus represents the percentage of the most common residue per column; higher bars denote greater conservation. Alignment quality represents the likelihood of observing mutations in a specific position+, based on BLOSUM 62 scores; a higher quality score indicates fewer mutations. Conservation represents the amino acid sequence similarity at specific regions of the protein. Conserved columns are marked with an asterisk (*) with a score of 11, while columns with a single mutation, where all properties were conserved, are marked with a ‘plus sign’ (+) with a score of 10. Columns with less conservation, evidenced by multiple mutations, are assigned conservation scores less than 10.

## Supplementary Tables (see separate Excel files)

**Table S2. Computational analysis of predicted functional domains in TiLV proteins.**

**Table S3. Six-frame translation of the TiLV genome.** Six-frame translation software (Bioline) was used to identify all potential peptide sequences from the TiLV genome (Israel isolate, Til-4-2011, accession nos. KU751814–KU751823), with a minimum length of 9 amino acids. The predicted primary ORF sequence of each segment is highlighted in yellow. Plus (+) and minus (-) indicate translation from positive and negative strand, respectively. Numbers after the (+) and (-) represent the frame and number of the *in silico* translation results. For instance, >TiLVS1+2.7 represents the seventh translational product of segment 1, derived from frame 2 of positive strand.

**Table S4. TiLV peptides identified by mass spectrometry.**

**Table S5. List of publicly available TiLV Segment 9 nucleotide sequences retrieved from the NCBI databases as of May 2025.** The country of origin and accession numbers are shown. Sequences highlighted in red and marked with red asterisk indicate additional exclusions from the S9-F3 analysis due to mutations that disrupt the predicted S9-F3 ORF.

**Table S6. Oligonucleotides used for the construction of GFP-tagged TiLV plasmids.**

**Table S7. Oligonucleotides used to introduce mutations into TiLV segment 9.**

